# Transcriptional Integration of Meiotic Prophase I Progression and Early Oocyte Differentiation

**DOI:** 10.1101/2025.01.06.631470

**Authors:** Kimberly M. Abt, Myles A. Bartholomew, Anna Nixon, Hanna E. Richman, Megan A. Gura, Kimberly A. Seymour, Richard N. Freiman

## Abstract

Female reproductive senescence results from the regulated depletion of a finite pool of oocytes called the ovarian reserve. This pool of oocytes is initially established during fetal development, but the oocytes that comprise it must remain quiescent for decades until they are activated during maturation in adulthood. In order for developmentally competent oocytes to populate the ovarian reserve they must successfully initiate both meiosis and oogenesis. As the factors that regulate the timing and fidelity of these early events remain elusive, we assessed the precise function and timing of the transcriptional regulator TAF4b during meiotic prophase I progression in mouse fetal oocytes. Compared to matched controls, E14.5 *Taf4b*-deficient oocytes enter meiosis I in a timely manner however, their subsequent progression through the pachytene-to-diplotene transition of meiotic prophase I is compromised. Moreover, this disruption of meiotic progression is associated with the reduced ability of *Taf4b*-deficient oocytes to repair double-strand DNA breaks. Transcriptional profiling of *Taf4b*-deficient oocytes reveals that between E16.5 and E18.5 these oocytes fail to coordinate the reduction of meiotic gene expression and the induction of oocyte differentiation genes. These studies reveal that TAF4b promotes the formation of the ovarian reserve in part by orchestrating the timely transition to meiosis I arrest and oocyte differentiation, which are often perceived as separate events.

## Introduction

The formation of a functional ovarian reserve during embryonic development is essential to female fertility in adulthood (Grive & Freiman, 2015). This finite pool of gametes will be continually depleted throughout a female’s reproductive lifespan until the onset of menopause, which occurs on average at 50±4 years of age. Onset of menopause before 40 years of age is considered premature, and this leads to infertility as well as an increased risk of mortality (van Noord et al., 1997). Premature menopause is clinically known as primary ovarian insufficiency (POI), but the cause of infertility cannot be determined in most cases. Individuals with POI display a diminished ovarian reserve measured by reduced serum anti-Mullerian hormone (AMH) levels (Nelson, 2009). Unfortunately, the timing and mechanisms of this fertility deficit remain unknown (Cordts et al., 2011; Cox & Liu, 2014; Nelson, 2009; Ruth et al., 2021) and little can be done toward its prevention or treatment. POI also disrupts key endocrine functions of the ovary, and this has detrimental effects on the health of extra-ovarian organs including the bones, heart, and brain (Rossetti et al., 2017). Defective ovarian reserve establishment is a likely contributor to the onset of POI, but the molecular mechanisms that govern this fundamental process are still being uncovered.

The initiation of meiosis and early oogenesis are two of the most critical processes that must occur during the establishment of the ovarian reserve. Oocytes are first formed during fetal development in cysts that breakdown producing individual primordial follicles consisting of a single dictyate-arrested oocyte surrounded by a flattened layer of pre-granulosa cells. These primary oocytes can be arrested for decades until after puberty when ovulation signals the completion of meiosis I and fertilization promotes the completion of meiosis II. The major developmental events of early oogenesis, such as cyst breakdown and fetal oocyte meiosis I arrest, are conserved between mice and humans. Thus, dissecting conserved genetic pathways underlying the embryonic development of the initial ovarian reserve between the mouse and human ovary is critical to develop better tools to detect a reduction of the ovarian reserve. These insights are key for devising novel strategies to preserve declining fertility and endocrine functions in individuals with POI.

TATA-box binding protein associated factor 4b (TAF4b) was originally identified as a gonadal-enriched subunit of the TFIID transcription initiation complex and a paralog of TAF4a in the mouse genome (Freiman et al., 2001). We more recently discovered TAF4b is critical for development of a healthy ovarian reserve in the mouse embryonic ovary (Freiman et al., 2001; Grive et al., 2014; Voronina et al., 2007). Importantly, adult female *Taf4b*-deficient mice model critical aspects of POI in women (Lovasco et al., 2010). Furthermore, TAF4b-dependent molecular events significantly overlap with those disrupted in Turner syndrome and Fragile X-associated POI (FX-POI), two prominent genetic examples of POI (Gura et al., 2022). In the mouse, we determined that *Taf4b* mRNA and protein are consistently enriched in the germ cells of the embryonic ovary compared to the somatic cells from embryonic day (E)11.5 to E18.5 (Gura et al., 2020). We further demonstrated that *Taf4b* mRNA expression significantly increases during meiotic differentiation, as do other alternative TFIID genes, including *Taf7l* and *Taf9b* (Gura et al., 2020). Most recently, we identified 570 TAF4b occupancy peaks using CUT&RUN in E16.5 mouse oocytes, of which 94% were located in the core promoter region just upstream of the transcription start site (TSS). Importantly, TAF4b peaks were identified in a large number of critical meiotic, oogenesis and POI-related gene promoters including *Fmr1* and *Dazl* (Gura et al., 2022).

It is still unknown if and how early meiotic events and oocyte differentiation are coordinated to establish a healthy ovarian reserve. A seminal study by Doshkin et al., in 2014 concludes that early meiotic prophase I and oocyte growth and differentiation are genetically separable events in the mouse (Doskin et al. 2014). As the timing of these early meiotic events and oocyte differentiation overlap, we sought to find a common regulator of both processes. Meiotic prophase I is a complex process that occurs asynchronously in the mouse ovary from E14.5-PND3 and can be broken down into five substages (leptonema, zygonema, pachynema, diplonema and dictyate) based on cytological appearance of the oocyte chromosomes (Cohen et al., 2015). Here we show that both *Taf4b* mRNA and protein are enriched during the pachytene stage of meiotic prophase I. While *Taf4b*-deficient oocytes are able to initiate meiosis I in a timely fashion, their subsequent progression is delayed and they largely fail to progress from pachynema to diplonema. Meiotic progression delays in the absence of TAF4b are coincident with increased RAD51 and gH2AX DNA damage foci suggesting that elevated levels of DNA damage underlie the excessive loss of these oocytes shortly after birth. Surprisingly, we observed that most *Taf4b*-deficient oocytes reached pachynema at E18.5 allowing us to compare gene expression of FACS-sorted control vs. *Taf4b*-deficient pachynema-enriched oocytes. At this timepoint, we discovered that *Taf4b*-deficient oocytes display increased expression of meiotic genes and decreased expression of oocyte differentiation genes compared to littermate controls. Together, these data suggest that TAF4b may coordinate the timely reduction of meiotic genes and upregulation of oogenesis genes required to establish a robust ovarian reserve. Although these two pathways are genetically separable, it is of great interest if a single transcriptional regulatory factor, TAF4b, simultaneously integrates both critical functions.

## Results

### TAF4b expression is enriched during pachynema in female germ cells

We previously found that *Taf4b* mRNA expression peaks at E16.5 and is expressed simultaneously with essential meiotic genes like *Hormad1*, *Meioc*, and *Dmc1* (Gura et al., 2022). These data led us to examine whether *Taf4b* is enriched in female germ cells in a particular substage of meiotic prophase I. To address this question, we integrated multiple publicly available single cell RNA-sequencing (scRNA-seq) datasets that together spanned ovarian reserve establishment in the mouse.

To perform the dataset integration, we reprocessed four datasets listed in **Figure 1A**, three of which performed scRNA-seq on whole ovaries and one of which performed scRNA-seq on Oct4-GFP^+^ oocytes (Ge et al., 2021; Niu & Spradling, 2022; J. J. Wang et al., 2020; Zhao et al., 2020). We first generated a merged Seurat object of all cells from each dataset. We then selected for high quality cells based on the following parameters: nFeature_RNA > 1000, nCount_RNA < 40000, and percent.mt < 20 (**Figure S1A-B**). We used Monocle3 to perform initial dimensionality reduction, and we found that the majority of Oct4-GFP^+^ oocytes from GSE130212 clustered with *Dazl* and *Ddx4*-positive germ cells obtained from the three datasets that performed scRNA-seq on whole ovaries (**Fig S1C, E-F**). We also found that germ cells isolated at similar timepoints from different datasets clustered together, suggesting the integration was successful (**Figure S1D**).

**Figure 1.**
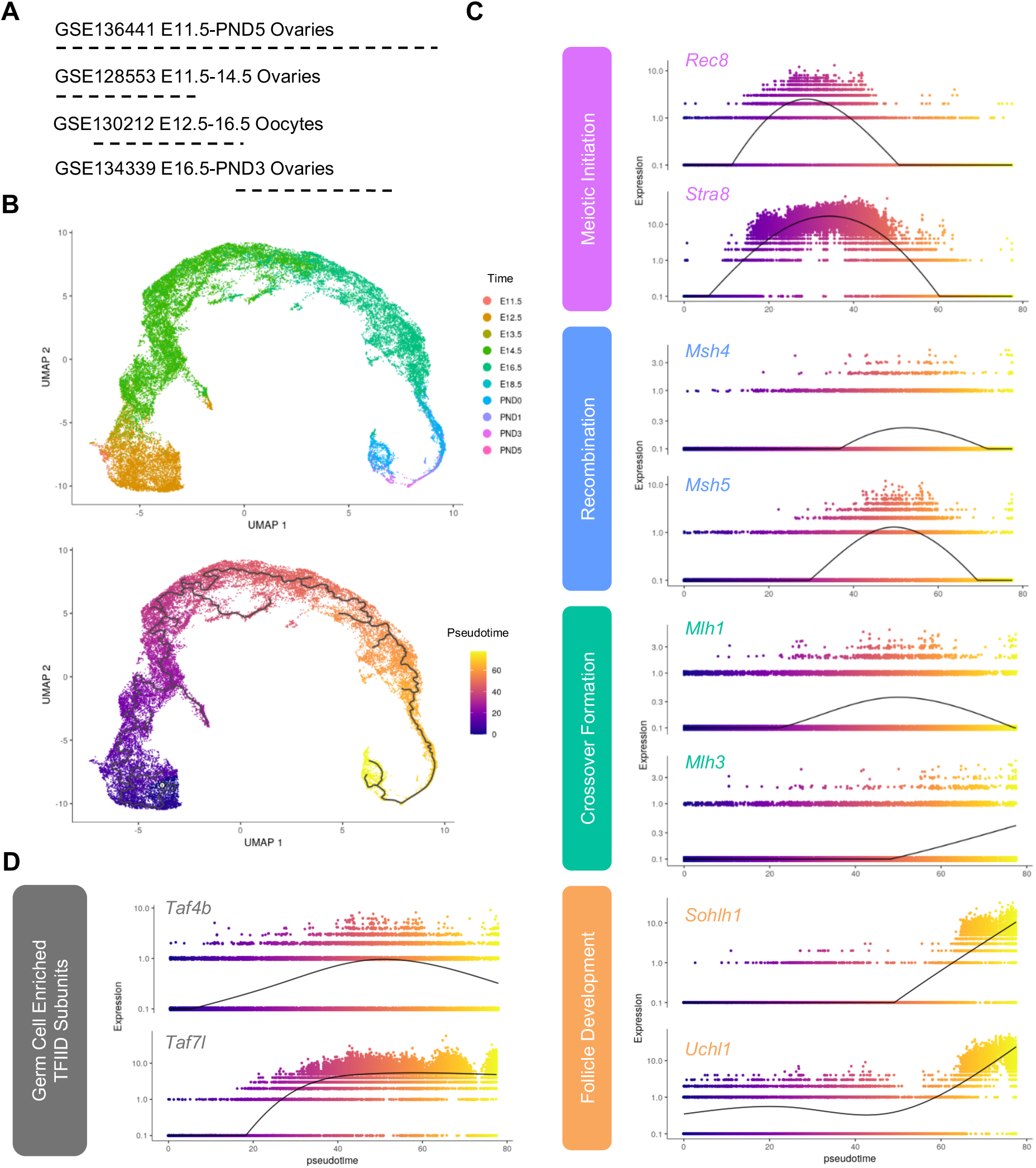
Integration of scRNA-seq data from oocytes during ovarian reserve establishment. (A) Schematic of datasets used for integration. (B) Uniform Manifold Approximation and Projection (UMAP) of female germ cells colored by time (upper panel) and pseudotime (lower panel). (C) Expression of candidate genes plotted in terms of pseudotime. Color of gene labels correspond to approximate prophase I stage, Figure 1 (D) Expression of germ-cell enriched TFIID components plotted in terms of pseudotime.

We found that the majority of *Ddx4* and *Dazl*-positive cells belonged to partition 1, and we used this subset of cells for downstream pseudotime analysis (**Figure S1G and 1B**). We found that our pseudotime analysis successfully recapitulated gene expression changes that occur throughout prophase I (**Figure 1C**). For example, expression of *Stra8*, which is a master regulator of meiotic initiation, peaks at the beginning of the pseudotime course (Anderson et al., 2008). *Rec8*, a cohesin protein that supports DNA replication prior to meiosis, peaks at a similar point along the pseudotime trajectory (Dokshin et al., 2013). Expression of genes required for recombination (*Msh4/5*) and crossover formation (*Mlh1/3*) peak after meiotic initiation (Bolcun-Filas & Handel, 2018). Furthermore, expression of oocyte enriched genes that are required for primordial follicle survival like *Sohlh1* and *Uchl1* peak at the end of the pseudotime trajectory (Niu & Spradling, 2022; Pangas et al., 2006; Woodman et al., 2022). We then evaluated the pseudotime trajectory of two germ cell enriched components of the TFIID complex: *Taf4b* and *Taf7l* (Gura et al., 2020). We found that expression of both increased at the section of the pseudotime course that was similarly enriched with zygotene and pachytene markers (**Figure 1D**).

To determine whether TAF4b protein is enriched in pachynema, we performed quantitative immunofluorescence on prophase I chromatin spreads isolated from E16.5 and E18.5 ovaries. Preparing chromatin spreads from fetal ovarian tissue allows us to both identify meiotic germ cells and characterize their substage based on the configuration of synaptonemal complex protein 3 (SYCP3) (Grive et al., 2016; Hwang et al., 2018). By staining chromatin spreads for TAF4b and DAPI in addition to SYCP3, we were able to use a recently published method to quantify the amount of TAF4b present relative to the DNA content of the cell in oocytes at specific stages of prophase I (Alexander et al., 2023). In E16.5 wildtype mice, we found oocytes in leptonema, zygonema, or pachynema and we observed that the staining pattern of TAF4b is diffuse throughout the chromatin spreads at each stage (**Figure 2A and C**). To validate our TAF4b antibody is specific in this assay, we performed immunofluorescence on prophase I spreads isolated from wildtype (*Taf4b+/+*) and *Taf4b*-deficient (*Taf4b-/-*) mice at E18.5, and we detected high levels of TAF4b signal in wildtype oocytes, but not in our *Taf4b-/-* spreads (**Figure S2**). By measuring fluorescence intensity of TAF4b in E16.5 chromatin spreads from wildtype mice, we found that its expression is significantly higher in pachytene oocytes even though it is present at all stages observed, which agrees with our findings from the scRNA-seq integration (**Figure 2B**). We performed a similar analysis in wildtype oocytes at E18.5, and we found that TAF4b expression is significantly higher in pachytene versus diplotene oocytes (**Figure 2C-D**). Together these data suggest that expression of *Taf4b* mRNA and protein are highest in pachytene oocytes during ovarian reserve establishment.

**Figure 2.**
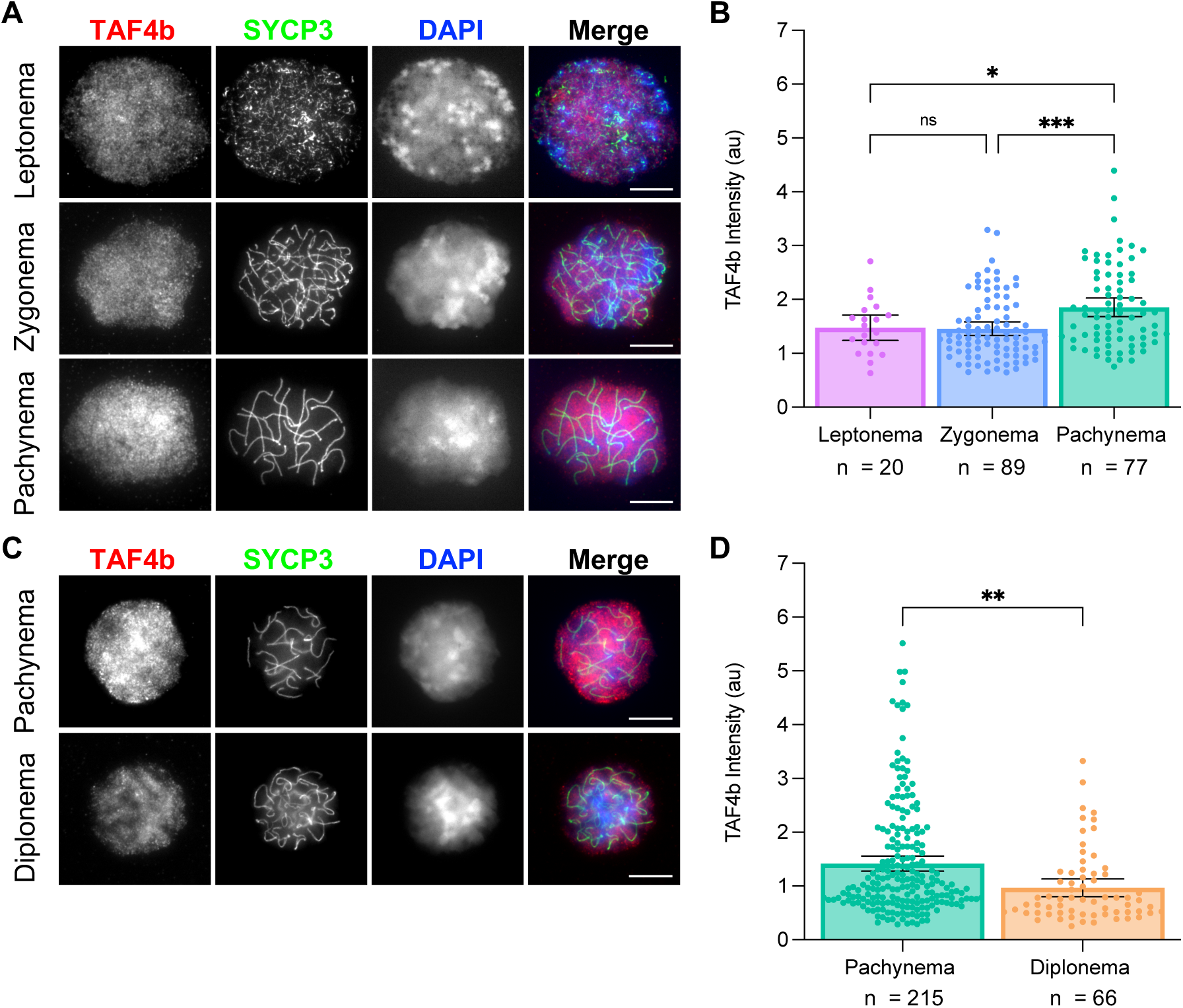
TAF4b is enriched in pachytene oocytes at E16.5 and E18.5. Prophase I chromatin spreads were prepared from wildtype ovaries at E16.5 and E18.5 using the drying down technique. Images of oocytes stained with TAF4b (red), SYCP3 (green), and DAPI (blue) at each prophase I stage present at E16.5 (A) or E18.5 (C), scalebar represents 10 μm. Quantification of TAF4b signal intensity in oocytes grouped by stage at E16.5 (B) or E18.5 (D), au= arbitrary units. Spreads were collected from 3 animals at E16.5 and 4 animals at E18.5, and n refers to the number of spreads used in analysis. Dots represent individual spreads, bar height represents the sample mean, and error bars represent standard error of the mean. Statistical significance for stage specific comparison at E16.5 was determined using an ordinary one-way ANOVA with multiple comparisons. For E18.5 samples a two-tailed T-test was used, ns=not significant, * p<0.05, ** p <0.01, and *** p <0.001.

### *Taf4b*-deficiency does not impede meiotic initiation at E14.5

Although *Taf4b* mRNA and protein expression are highest during pachynema, it is still expressed at notable levels during earlier stages of prophase I. Moreover, previous work has shown that expression of *Taf4b* is modulated by the regulator of meiotic initiation *Stra8* (Anderson et al., 2008; Gura et al., 2020). This led us to investigate whether initiation of meiosis I is perturbed in the absence of *Taf4b*. Fetal ovary tissue sections were isolated from E14.5 *Taf4b+/+* and *Taf4b-/-* mice and stained with antibodies for TRA98, as a germ cell marker, and STRA8, as a marker of meiotic initiation. We counted the number of TRA98 or STRA8-positive cells per section and divided this number by the DAPI-positive area of the ovarian tissue to measure the density of total and meiotic germ cells in ovaries from each genotype. We found that *Taf4b+/+* and *Taf4b-/-* ovaries were largely indistinguishable from each other at E14.5 (**Figure 3A-B**). We found no significant difference in the density of TRA98-positive germ cells or STRA8-positive meiotic germ cells between *Taf4b+/+* versus *Taf4b-/-* mice (**Figure 3C-D**). These findings are consistent with our previous bulk RNA-seq analysis of E14.5 Oct4-GFP^+^ oocytes isolated from *Taf4b+/+* and *Taf4b-/-* ovaries, which found little to no differential gene expression between the two genotypes at this timepoint (Gura et al., 2022).

**Figure 3.**
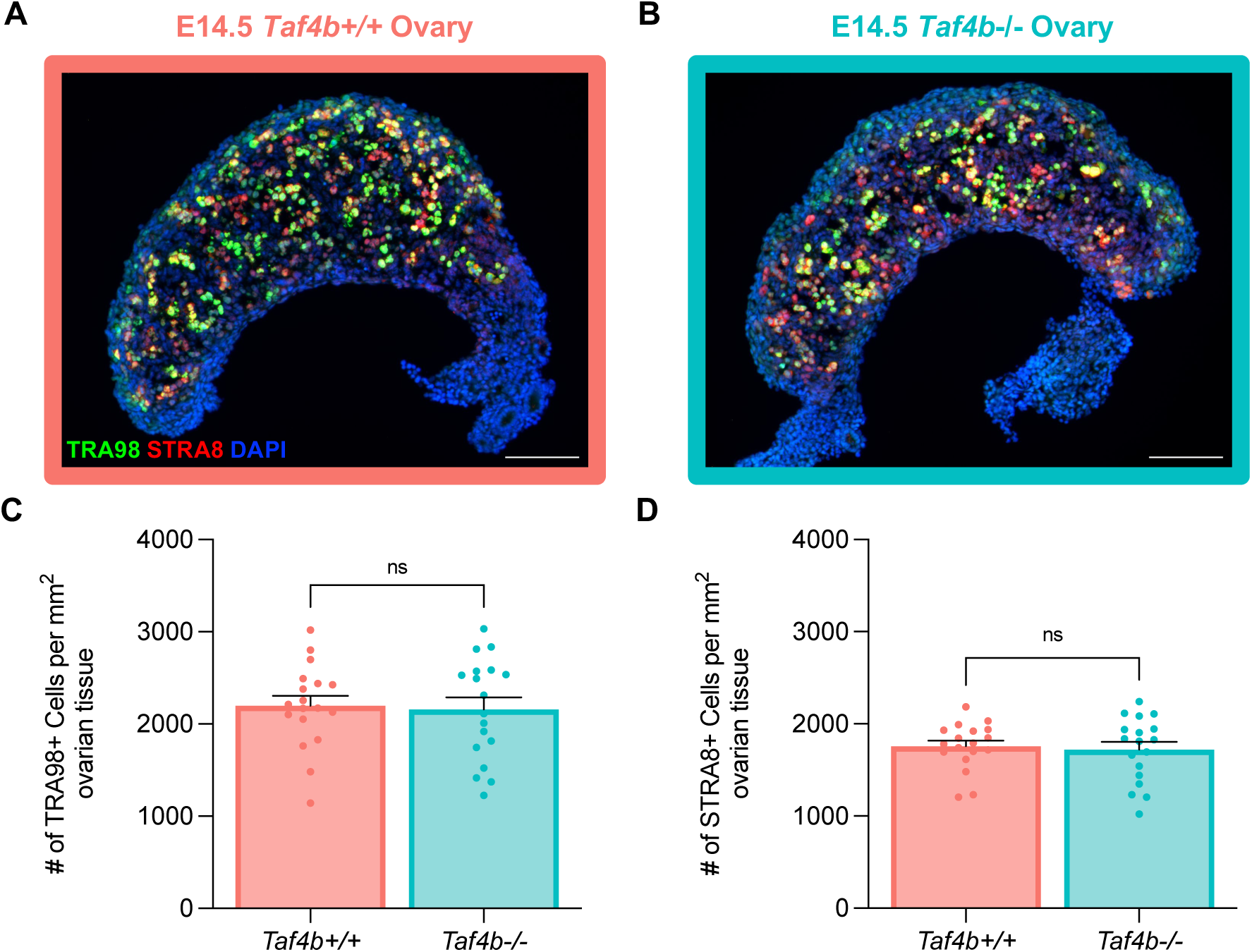
*Taf4b-/-* ovaries initiate meiosis normally at E14.5. Images of E14.5 *Taf4b+/+* (A) and *Taf4b-/-* (B) ovary sections stained with TRA98 (green) as a marker of total germ cells, STRA8 (red) as a marker of meiotic germ cells, and DAPI (blue) as a marker of cell nuclei. Scalebar represents 100 μm. Quantification of TRA98 (C) or STRA8 (D) cell density of tissue sections per genotype. Cell density was calculated by dividing the number of positively marked cells by the area of the tissue section. Sections were collected from three E14.5 ovary pairs per genotype. Dots represent individual sections, bar height represents the sample mean, and error bars represent standard error of the mean. Statistical significance for genotype comparisons was determined using a two-tailed T-test, ns=not significant.

### *Taf4b*-deficient oocytes experience delayed meiotic prophase I progression

While fetal oocytes can initiate meiosis properly in the absence of *Taf4b*, *Taf4b-/-* oocytes are unable to reach pachytene as efficiently as their wildtype counterparts at E16.5 and the vast majority of *Taf4b-/-* oocytes undergo apoptotic cell death shortly after birth (Grive et al., 2014). These data led us to further examine the precise timing of prophase I progression in *Taf4b-/-* oocytes just prior to and immediately after birth. To accomplish this, we made chromatin spreads from three *Taf4b+/+* and *Taf4b-/-* mice at E16.5, E18.5, as well as PND0 and stained for SYCP3 to determine the proportion of oocytes in each substage of prophase I. We isolated 50-100 spreads per animal and then performed a Chi-square test to determine whether there was a significant difference in the percentage of oocytes at each stage between the two genotypes. Our analysis of E16.5 *Taf4b-/-* spreads corroborated our previous findings that these oocytes do not reach pachytene as efficiently as wildtype controls (**Figure 4A**) (Grive et al., 2016). Intriguingly, the majority of *Taf4b-/-* oocytes reach pachynema by E18.5, but roughly 20% of *Taf4b-/-* germ cells remain in zygonema whereas a similar percentage of *Taf4b+/+* oocytes have proceeded to diplonema (**Figure 4A**). By PND0 most *Taf4b+/+* oocytes are in diplonema, but 80% of *Taf4b-/-* germ cells remain in pachynema (**Figure 4A**). Together these data indicate that *Taf4b-/-* oocytes are unable to progress through prophase I in a timely manner and are largely arrested in pachynema at PND0.

**Figure 4.**
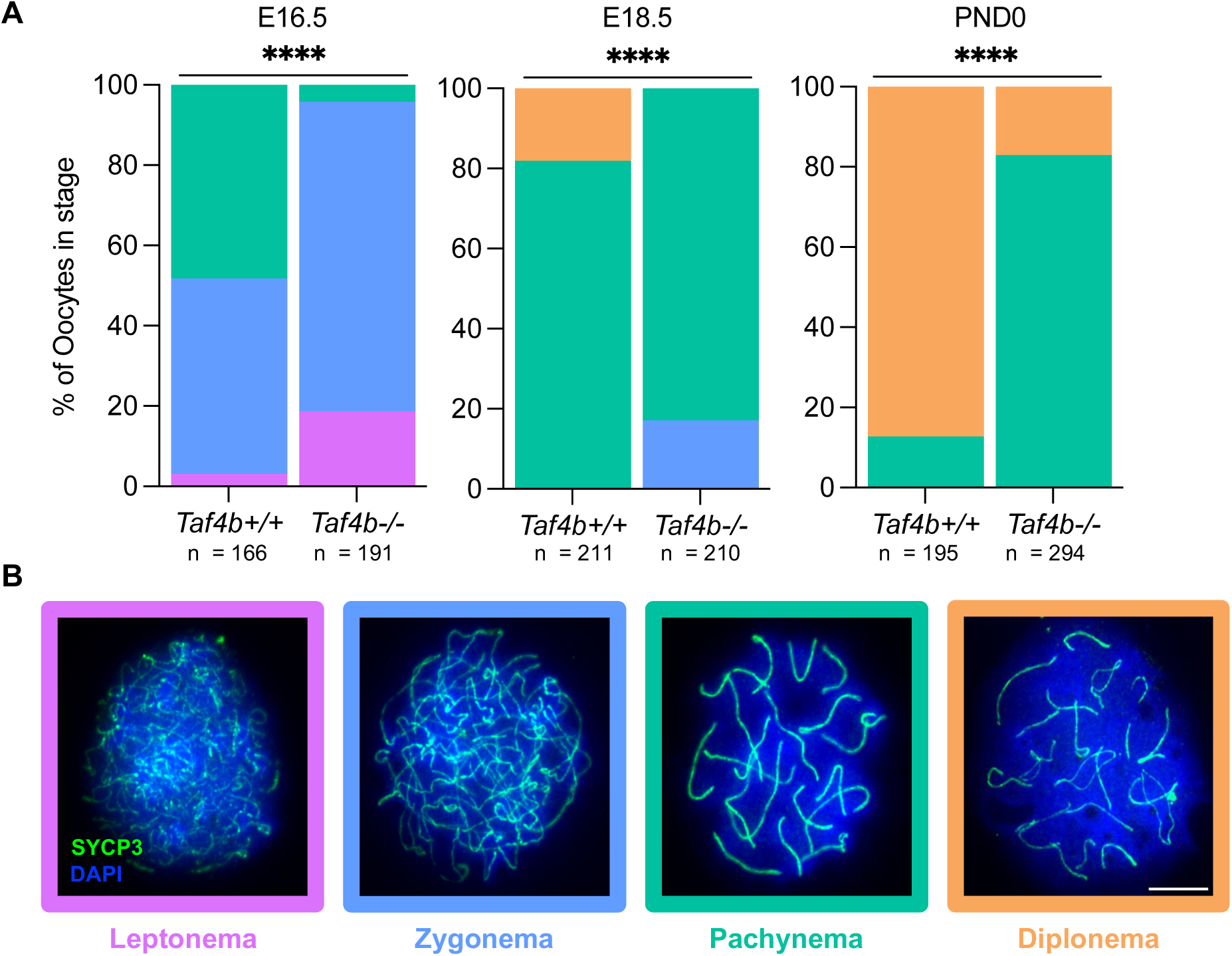
*Taf4b-/-* ovaries experience delays in prophase I progression at E16.5, E18.5 and PND0. (A) Proportion of *Taf4b+/+* and *Taf4b-/-* chromatin spreads found at each substage of prophase I at E16.5, E18.5, and PND0. Substage was determined by spatial configuration of SYCP3 and representative images of spreads at each stage are shown in (B). Spreads were stained with SYCP3 (green) and DAPI (blue) and scalebar represents 10 μm. Color of labels in stacked bar chart corresponds to prophase I stage, leptonema (purple), zygonema (blue), pachynema (teal), diplonema (orange). Spreads were collected from three animals per genotype at each timepoint and n refers to the number of spreads analyzed per stage. Statistical significance was calculated using a Chi-square test, **** p<0.0001.

### *Taf4b-*/- oocytes have excessive DNA damage during pachynema and diplonema

During fetal development and early postnatal life there is a significant amount of fetal oocyte attrition (FOA) in the mammalian ovary, likely driven by quality control processes that eliminate defective germ cells and consequently ensure the survival of highest quality oocytes (Grive, 2020; Grive & Freiman, 2015; Hunter, 2017). Errors in prophase I such as persistent DNA damage and chromosomal asynapsis are known triggers of excessive FOA (Di Giacomo et al., 2005; Huang & Roig, 2023; Ravindranathan et al., 2022; Rinaldi et al., 2017). We have previously shown that *Taf4b*-/- oocytes have increased levels of asynapsis at E16.5, and more recent work has demonstrated that *Atr* as well as other genes involved in DNA repair are differentially expressed in E16.5 *Taf4b*-deficient oocytes (Grive et al., 2016; Gura et al., 2022). To determine whether *Taf4b-/-* oocytes have excessive DNA damage, we stained prophase I spreads from *Taf4b+/+* and *Taf4b-/-* with antibody against γH2AX as a marker of unrepaired double strand breaks (DSBs). To complete this analysis, we pooled spreads from three timepoints (E16.5, E18.5, PND0) and separated spreads by substage in order to compare levels of γH2AX between *Taf4b+/+* and *Taf4b-/-* oocytes using quantitative immunofluorescence. We found that *Taf4b+/+* and *Taf4b-/-* oocytes had similar levels of γH2AX signal during zygonema; however, *Taf4b-/-* oocytes have elevated levels of γH2AX signal during pachynema and diplonema (**Figure 5A-F**). We did not compare levels of γH2AX signal between *Taf4b+/+* and *Taf4b-/-* oocytes in leptonema because there are not sufficient numbers of *Taf4b+/+* oocytes in this stage at the timepoints analyzed. We also quantified the number of RAD51 foci in *Taf4b+/+* and *Taf4b-/-* oocytes at E16.5 to evaluate single strand invasion during homologous recombination. We found a similar number of RAD51 foci in *Taf4b+/+* and *Taf4b-/-* oocytes during leptonema as well as zygonema (**Figure S3A-D**). However, in *Taf4b-/-* pachytene oocytes, we observed an increased number of RAD51 foci compared to *Taf4b+/+* (**Figure S3E-F**). These data suggest that levels of DNA damage are normal in *Taf4b-/-* oocytes during the initial stages of prophase I, but that they remain elevated and persistent during pachynema and diplonema, when wild-type oocytes are completing DNA repair. This is consistent with our findings that *Taf4b-/-* oocytes largely initiate prophase I similarly to controls but they are not able to progress to later stages in a timely manner.

**Figure 5.**
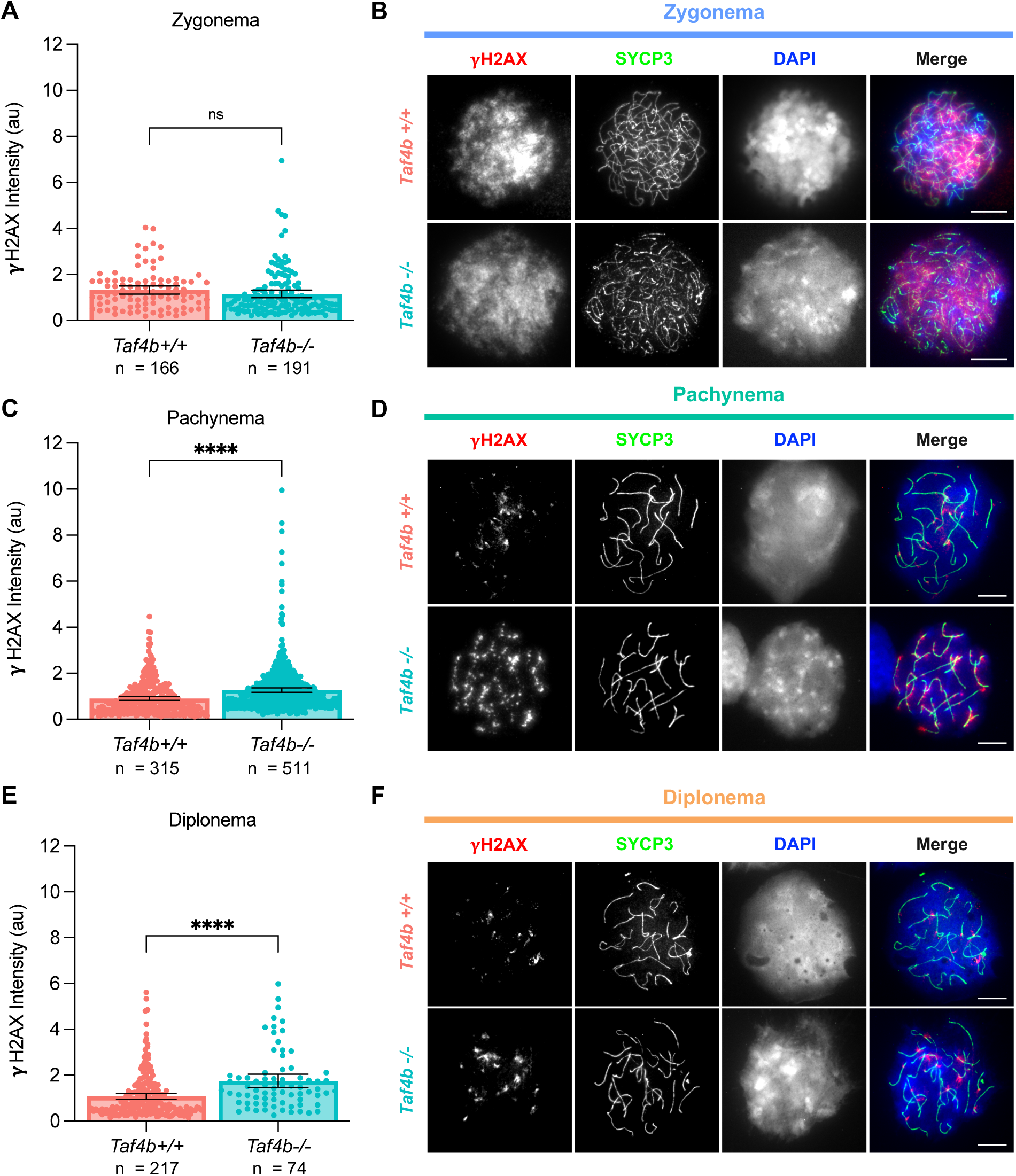
*Taf4b-/-* oocytes have elevated levels of γH2AX during pachynema and diplonema. Quantification of γH2AX signal intensity in chromatin spreads during zygonema (A) pachynema (C) and diplonema (E). Spreads were pooled from three E16.5, E18.5, and PND0 mice per genotype, n = the number of spreads analyzed, au = arbitrary units. Images of spreads stained with γH2AX (red) SYCP3 (green) and DAPI (blue) from each genotype during zygonema (B) pachynema (D) and diplonema (F). Scalebar represents 10 μm. Dots in graphs represent individual spreads, bar height represents the sample mean, and error bars represent standard error of the mean. Statistical significance for stage specific comparisons was determined using a two-tailed T-test, ns=not significant, and **** p <0.0001.

### TAF4b promotes the pachytene-diplotene transition in E18.5 oocytes

To understand the transcriptome-level changes in *Taf4b*-/- oocytes, we previously performed bulk RNA-sequencing on Oct4-GFP^+^ germ cells isolated from control and *Taf4b-/-* ovaries at both E14.5 and E16.5. We found few transcriptional changes at E14.5, which agrees with these findings that meiotic initiation occurs normally in the absence of *Taf4b* (**Figure 3**). However, we found a significant number of differentially expressed genes at E16.5; for example, essential oogenesis genes *Sohlh1* and *Nobox* were reduced in *Taf4b-/-* oocytes (Gura et al., 2022). In addition, we observed reduced expression of X chromosome genes and that *Taf4b-/-* oocytes share significant transcriptome similarities with fetal oocytes collected from a mouse model of Turner Syndrome (Gura et al., 2022). Surprisingly, we did not observe significant changes in expression of meiotic genes at E16.5 (Gura et al., 2022). We hypothesized that this could have been caused by the heterogeneity of the Oct4-EGFP^+^ germ cell population present at E16.5 when *Taf4b-/-* oocytes are largely in leptonema and zygonema whereas controls have begun to reach pachynema (**Figure 4**). As *Taf4b* mRNA expression is enriched in pachynema and *Taf4b-/-* oocytes remain in this stage for an extended period of time before they are culled, it is essential to understand what genes are regulated by TAF4b at this meiotic sub-stage.

To accomplish this goal, we performed bulk RNA-seq on sorted Oct4-EGFP^+^ oocytes from five *Taf4b* +/+, *Taf4b* +/-, and *Taf4b* -/- ovary pairs at E18.5 (cell numbers for each RNA-seq sample can be found in **Figure S4**). Although *Taf4b-/-* oocytes still display a lag in prophase I progression at E18.5, both *Taf4b+/+* and *Taf4b-/-* oocytes are largely in pachynema (**Figure 4**). We included *Taf4b+/-* oocytes in addition to *Taf4b+/+* oocytes as controls because we found that *Taf4b+/-* fetal oocytes have a slight delay in prophase I progression even though the adults are fertile (data not shown). The resulting principal component analysis (PCA) plot shows the *Taf4b*-/- samples clustering together away from both *Taf4b+/+* and *+/-* control samples (**Figure 6A**). We identified 1551 differentially expressed genes (DEGs) between *Taf4b*+/+ or *Taf4b+/-* and *Taf4b-/-* oocytes, which were defined as protein-coding, average transcripts per million (TPM) > 1, padj < 0.05, and log2FoldChange > l0.6l (**Fig 6B**, **Table S1**). From this list of DEGs, 1070 were decreased (downregulated) and the remaining 481 were increased (upregulated). As elevated DNA damage and asynapsis results in transcriptional silencing, the higher number of downregulated genes is consistent with our finding that DNA damage is elevated in *Taf4b-/-* oocytes (**Figure 5** and **S3**) (Cloutier et al., 2015). Moreover, if TAF4b acts solely as a transcriptional activator would we expect more genes to be downregulated in its absence. Strikingly, *Uchl1* and *Sohlh1* were two of the most significantly downregulated genes (**Figure S5B**). We also found other well characterized oogenesis genes like *Nobox, Figla, Kit, Lhx8, Foxo3,* and *Zp3* were downregulated in E18.5 *Taf4b-/-* oocytes (Choi et al., 2008; Jones & Pepling, 2013; Rajkovic et al., 2004; Shimamoto et al., 2019; Soyal et al., 2000; Suzumori et al., 2002; Z. Wang et al., 2020). Finding *Nobox* and *Figla* as DEGs corroborates our previous research that TAF4b directly binds to the promoter regions of these genes in E18.5 wildtype ovaries (Grive et al., 2016). In addition, H2AX was one of the most significantly upregulated genes, and we have shown its phosphorylated protein product (γH2AX) is elevated in *Taf4b-/-* oocytes during pachynema and diplonema (**Figure 5**).

**Figure 6.**
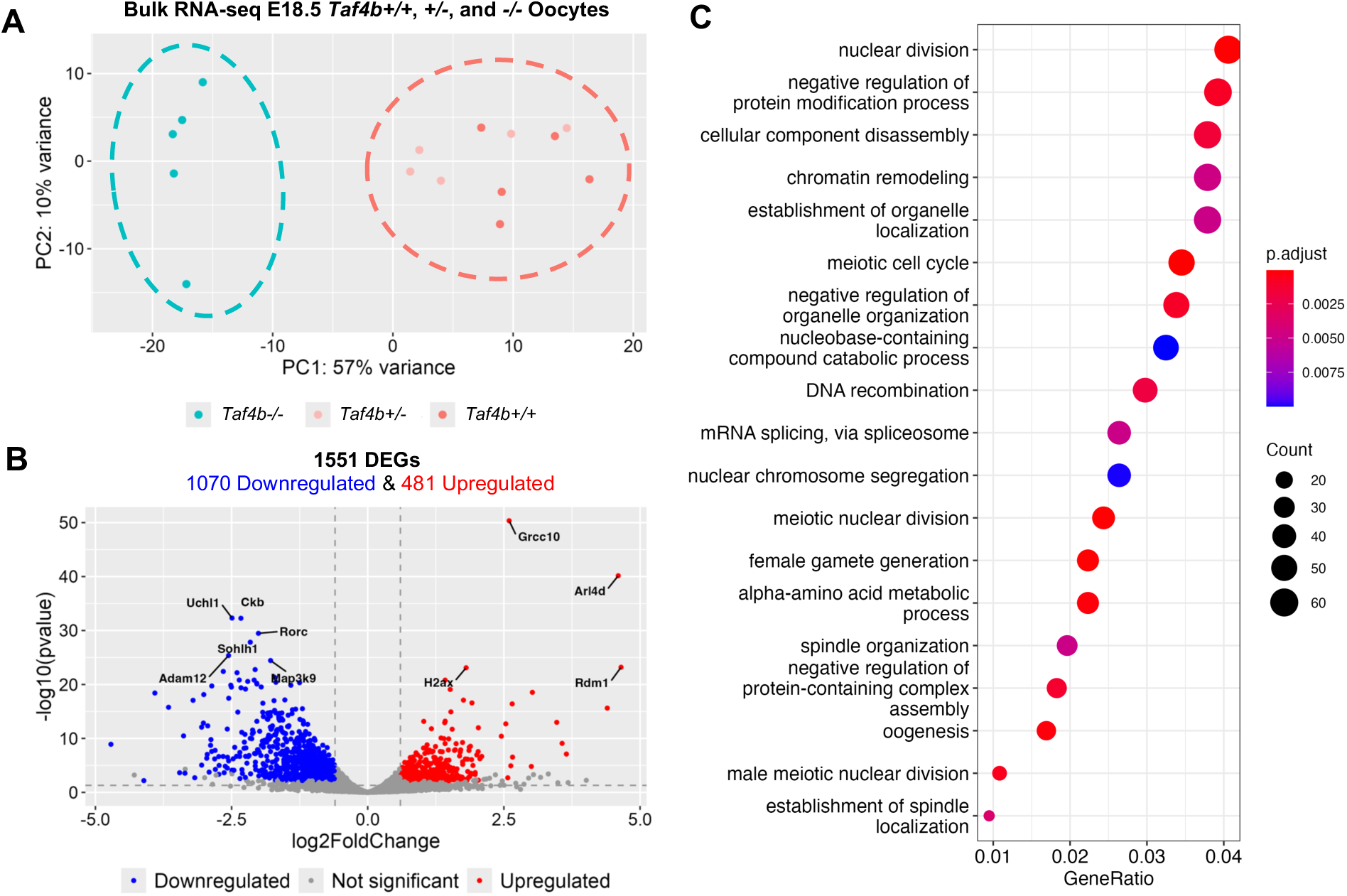
Bulk RNA-seq of E18.5 *Taf4b+/+, +/-,* and *-/-* oocytes. (A) PCA plot of E18.5 samples labelled based on genotype. Dashed circles represent samples compared in differential gene expression analysis. (B) Volcano plot of differentially expressed genes (protein-coding, padj <0.05, avg TPM > 1, log2FoldChange > l0.6l). The top 10 most significant DEGs are labelled. Dashed lines represent padj and log2FoldChange cutoffs. (C) Dotplot of GO biological process analysis of all 1551 DEGs.

To gain a comprehensive understanding of how gene expression is perturbed in E18.5 *Taf4b-/-* oocytes, we performed gene ontology (GO) analysis of all 1551 DEGs (**Figure 6C**). Multiple GO categories associated with oogenesis and meiosis were enriched in our DEGs allowing us to prepare heatmaps of genes lists associated with each process to examine how their expression was altered in *Taf4b-/-* oocytes (**Figure 7A-B** and **S6**). We found that genes involved in meiotic recombination (*Spo11*, *Dmc1*, *Meiob*, *Msh4*, *Brme1*, *Hormad1*, *M1ap*, and *Mcmdc2*) and meiosis-specific telomere movements (*Spdya*, *Majin*) were upregulated in E18.5 *Taf4b-/-* oocytes compared to controls (**Figure 7AD**) (Bolcun-Filas & Handel, 2018). Expression of these genes could be elevated in *Taf4b-/-* oocytes because they may be in an earlier state of pachytene than controls. Interestingly, recent work has shown that *Meioc* is required for the extension of prophase I, and it is possible that upregulation of this gene at E18.5 in *Taf4b-/-* oocytes plays a role in the observed prophase I lag (Abby et al., 2016; Soh et al., 2017). We also found some genes involved in meiosis and DNA recombination were downregulated (*Mnd1*, *Mus81*, and *Ercc1*) in the absence of *Taf4b*. Dysregulation of genes involved in DNA recombination is particularly interesting given that pachytene *Taf4b-/-* oocytes have reduced numbers of MLH1 foci and fail to properly form crossovers at E18.5 (Grive et al., 2016). When looking at DEGs related to oogenesis and female gamete generation, we noticed that most of the upregulated genes were also associated with meiosis (*Hormad1*, *Spo11*, *Spdya*, *Meioc*, and *Mcmdc2*), whereas genes associated with oocyte differentiation were downregulated (*Nobox, Figla*, *Sohlh1*, *Foxo3*, and *Zp3*) (**Figure 7B**). Additional factors important for early oocyte development like *Uchl1*, *Kit*, *Lhx8*, and *Trp63* are similarly downregulated in *Taf4b-/-* oocytes (**Figure 6B, Table S1**). Furthermore, we compared our E18.5 *Taf4b-/-* DEGs with a list of nearly 200 genes found to be enriched in diplotene/dictyate oocytes using scRNA-seq, and the majority of these genes were downregulated in the absence of Taf4b (**Figure S5A-B**). This is consistent with our findings that few *Taf4b-/-* oocytes are in diplonema at E18.5 and that TAF4b may play a role in regulating expression of genes required to reach diplonema. Overall, E18.5 *Taf4b-/-* oocytes fail to properly express correct levels of meiotic genes and are unable to activate expression of genes required for oocyte differentiation and growth.

**Figure 7.**
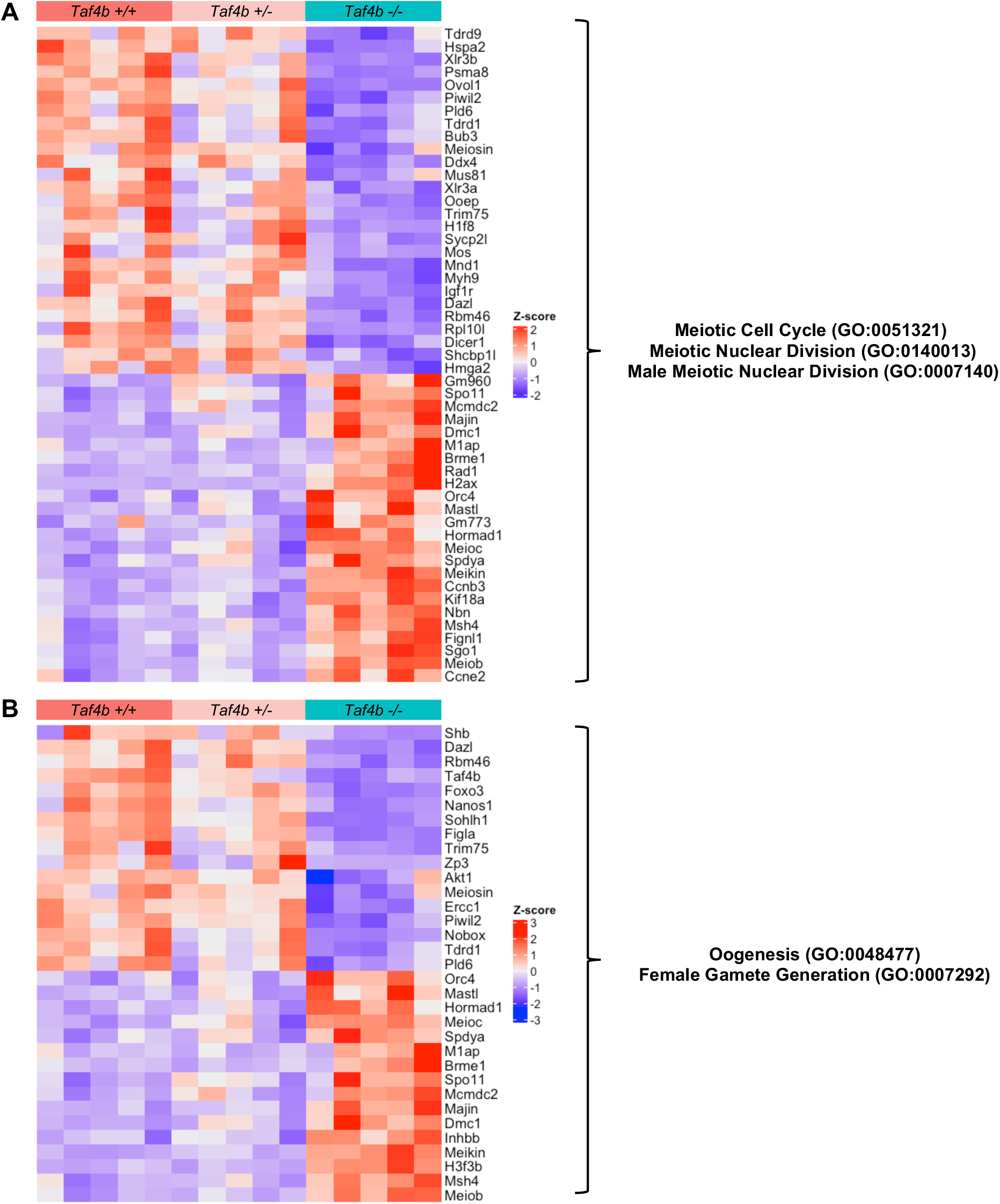
Dysregulation of genes involved in meiosis and oogenesis in E18.5 *Taf4b-/-* oocytes. (A) Heatmap of genes in meiotic cell cycle, meiotic nuclear division, or male meiotic nuclear division GO categories that were differentially expressed in E18.5 *Taf4b-/-* oocytes. (B) Heatmap of genes in oogenesis or female gamete generation GO categories that were differentially expressed in E18.5 *Taf4b-/-* oocytes. Heatmaps were generated using based normalized counts output from DESeq2.

To understand how gene expression is perturbed in the absence of TAF4b over time, we compared our bulk RNA-seq data from E16.5 (Gura et al., 2022) and E18.5 (this study) *Taf4b-/-* oocytes. We reprocessed our E16.5 *Taf4b+/-* and *-/-* samples to account for updates made to software packages used in our computational pipeline as well as updated gene annotations in the GO database. We found a slightly lower number of DEGs than we did previously (951 versus 964) but these DEG lists were over 99% identical (**Figure S6A-B**, **Table S2**). Intriguingly, chromatin remodeling was one of the top GO categories present in our E16.5 and E18.5 *Taf4b-/-* DEGs (**Figure 6C and S6C**). Genes present in the chromatin remodeling category differ between with E16.5 and E18.5 *Taf4b-/-* DEG lists, but we did find that the H3K4 demethylase *Kdm1b* was consistently downregulated. We also found that several histones (*H1f3*, *H1f4*, *H1f6*, *H2ac1*, *H2bc1*, *H2bc22*, *H2bc6*, *H4c8*, *H4c9*) were upregulated at both timepoints but their expression is still very low overall. Unfortunately, little is known about the epigenetic landscape of mouse oocytes at E16.5 and E18.5, but these data suggest that changes in chromatin structure are important for female germ cell development at this stage (Hu et al., 2023). We also found that genes associated with reproductive development like *Ddx4*, *Nobox*, and *Sohlh1* were consistently downregulated in E16.5 and E18.5 *Taf4b-/-* oocytes. Moreover, we found that expression of the X chromosome is also reduced at E18.5, and that there is a significant overlap between *Taf4b-/-* and XO DEGs at this timepoint (**Figure S7A-C**). GO analysis of the 642 overlapping DEGs revealed that there was shared disruption of genes involved in meiotic cell cycle, chromatin remodeling, and oogenesis (**Figure S7D**). Together, these data suggest that *Taf4b-/-* oocytes consistently fail to increase expression of genes required for early oocyte development at both E16.5 and E18.5.

## Discussion

From their origins as mitotic primordial germ cells (PGCs) in the fetal ovary, female germ cells undergo a multitude of precise developmental transitions to achieve meiotic and fertilization competence (Grive & Freiman, 2015). In the early mammalian embryo, Bone Morphogenetic Protein (BMP) signals from extraembryonic mesoderm signal PGC specification via the expression of three transcription factors PRDM1, PRDM14 and TFAP2C (Saitou & Yamaji, 2012). After migration to the bipotential gonad, sex-specification of the supporting somatic cells into pre-granulosa cells leads to retinoic acid-dependent STRA8 and retinoic acid-independent ZGLP1 expression which are both germ cell-specific transcription regulators of the transition from mitosis to meiosis (Anderson et al., 2008; Ishiguro, 2023; Miyauchi et al., 2017; Miyauchi et al., 2018; Nagaoka et al., 2020). However, transcription factors that regulate the timing of progression through meiosis I, early oocyte differentiation and survival required for proper dictyate arrest remain unknown. Moreover, whether these simultaneous but separable genetic processes have common regulators is unknown. Here, using several genomic and developmental approaches, we find that although TAF4b is not required for meiotic initiation at E14.5, it is essential for the timely progression of meiotic prophase I and its absence largely results in pachytene-arrested oocytes. At the genomic level, TAF4b-deficient pachytene meiocytes display elevated double-strand DNA breaks, fail to turn down early meiotic gene expression and turn up oocyte differentiation gene expression. Delayed and then stalled meiotic prophase I progression likely leads excessive oocyte attrition at birth and POI-like phenotypes in adult female mice (**Figure 8**).

**Figure 8.**
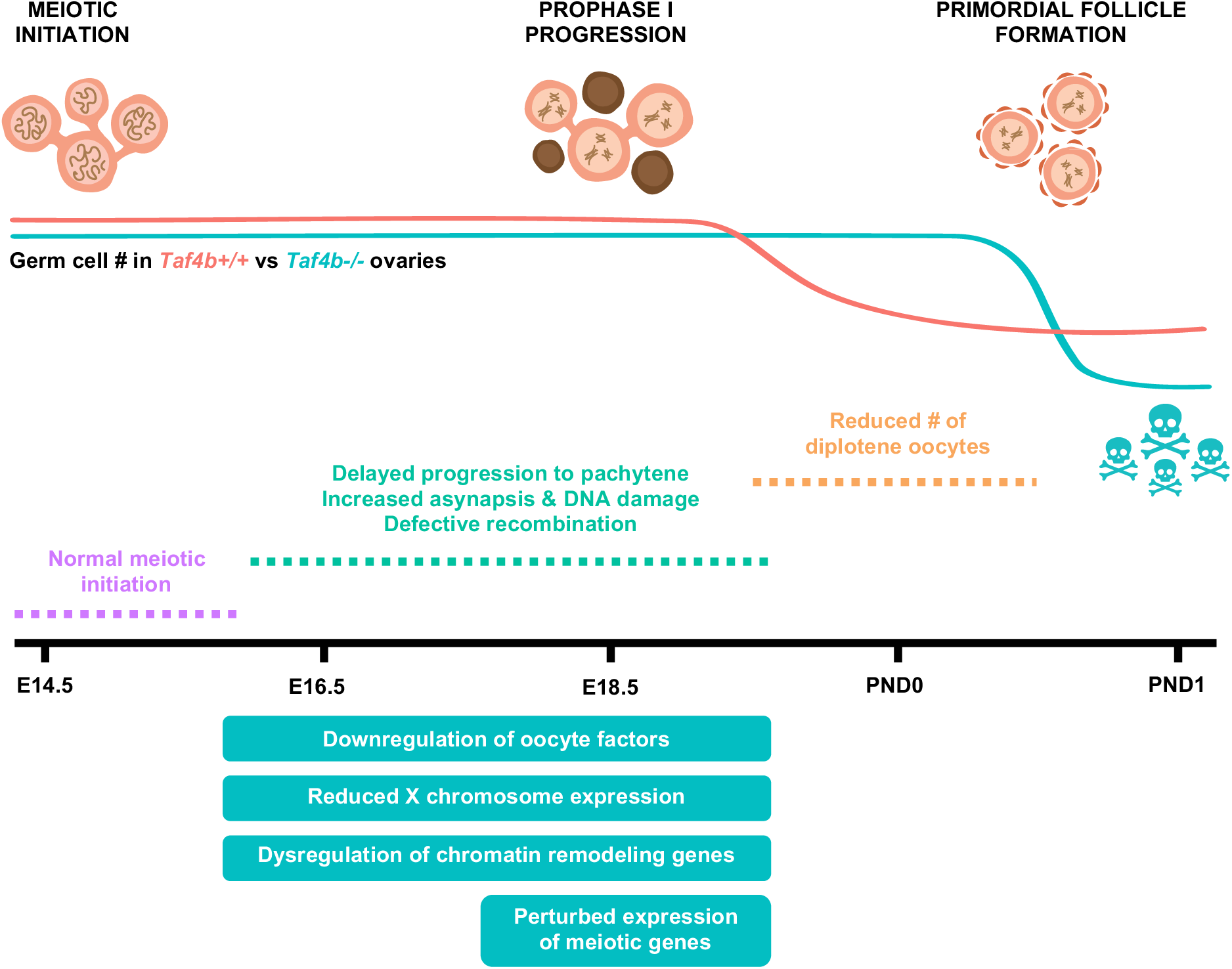
Model of defective ovarian reserve establishment in *Taf4b-/-* mice. Schematic timeline of major events that occur during ovarian reserve establishment in mice (top). Estimations of oocyte numbers in *Taf4b+/+* (red line) and *Taf4b-/-* (blue line) mice throughout time course are depicted below schematic. In the absence of TAF4b, there is a reduced number of oocytes at PND1. *Taf4b-/-* oocytes at E18.5 and PND0 are unable efficiently reach diplotene and are largely stuck in pachytene. *Taf4b-/-* oocytes do not reach pachytene in a timely manner at E16.5 but initiation of meiosis occurs normally at E14.5. At E16.5 and E18.5 *Taf4b-/-* oocytes downregulate genes essential for oocyte development, display reduced expression of the X chromosome, and dysregulate chromatin remodeling genes. In addition, expression of genes required for successful completion of prophase is highly perturbed in *Taf4b-/-* oocytes at E18.5.

Although the fetal human ovary is not easily amenable to such functional studies, there are a number of important reasons to conclude that mouse TAF4b functions are conserved during human oogenesis. First, the dynamic expression of *TAF4b* mRNA in human fetal oocytes is consistent with what we observe in the mouse (Gura et al., 2020). Second, several human genetic studies linked TAF4b expression to POI and human oocyte quality (Di Pietro et al., 2008; Knauff et al., 2009). Recently, a GWAS study of human age at natural menopause demonstrated 290 potential POI-related genes many of which are mis-expressed in *Taf4b*-deficient mouse E18.5 embryonic oocytes (Ruth et al., 2021). Finally, a heterozygous *TAF4b* mutation is predicted in the genome of the mother of three infertile brothers that display homozygous and truncating mutations in *TAF4b* causing their non-obstructive azoospermia (Ayhan et al., 2014). Together, our data strongly support that TAF4b plays similar roles during mouse and human gametogenesis. In this study, we have identified that TAF4b functions to promote meiosis I progression through the pachytene-to-diplotene transition in late embryonic mouse oocytes. In addition, we identified many critical meiotic and oocyte differentiation genes that are increased and decreased, respectively, in *Taf4b-*deficient E18.5 oocytes compared to controls. We infer from these data that TAF4b both directly and indirectly regulates the expression of these gene programs which are required for proper homologous recombination, oocyte differentiation and dictyate arrest. However, the precise mechanism of TAF4b and its associated partners to coordinate the integration of all these events is still unknown. Recently, we used CUT&RUN to map its occupancy to the core promoter regions just upstream of the TSS of several hundred E16.5 oocyte genes (Gura et al., 2022). These data are consistent with TAF4b participating in core promoter recognition as part of an embryonic and germ-cell specific form of TFIID. Our data also reflects the exciting possibility that a TAF4b-containing transcription complexes likely fine tunes transcription levels (up or down) rather than turning genes on or off. This buffering function is likely important to move oocytes through delicate meiotic transitions with maximum developmental fidelity. Oocyte aneuploidy largely results from meiotic prophase I errors and TAF4b is required for orchestrating the proper timing through pachynema. In the absence of this regulation by TAF4b, meiotic prophase I progression is largely halted at pachynema and the majority of such error-prone oocytes are culled from the ovary by excessive oocyte attrition soon after birth.

Finally, several additional levels of gene regulation are known and likely to be coordinated during these early meiotic and oocyte differentiation events. These include chromatin remodeling and the post-transcriptional and post-translational regulation of RNA and proteins, respectively. TAF4b may help to integrate several diverse aspects of oocyte gene expression. This is most notable in the GO categories revealed to be required during the transition from meiosis to oogenesis, **Figure 6**, like protein modification, chromatin remodeling and RNA splicing. In addition, many sequence-specific transcription factors like NOBOX, FIGLA, SOHLH1, and FOXO3 are dysregulated in the absence of TAF4b and likely have their own consequences on gene transcription. UCHL1, a potent regulator of ubiquitin-mediated proteolytic events in early oocytes is also reduced in expression (Woodman et al., 2022). Finally, MEIOC, a post-transcriptional regulator of meiotic prophase I progression displays reduced expression in TAF4b-deficient oocytes (Soh et al., 2017). It is likely that these developmental transitions are so delicate and dynamic that it is the concerted integration of gene expression at all these levels, including TAF4b, that ensures that developing oocytes correctly traverse these early meiotic and oocyte differentiation programs in a timely fashion.

## Materials and Methods

### Ethics statement

This study was approved by Brown University IACUC protocol #21-02-0005. The primary method of euthanasia is CO_2_ inhalation and the secondary method used is cervical dislocation both as per American Veterinary Medical Association (AVMA) guidelines on euthanasia.

### Mice

Mice that were homozygous for an Oct4-EGFP transgene (The Jackson Laboratory: B6;129S4-*Pou5f1^tm2Jae^*/J) were backcrossed to the C57BL/6 line and mated for collections used to prepare meiotic spreads. Mice that were homozygous for an Oct4-EGFP transgene (The Jackson Laboratory: B6;129S4-*Pou5f1^tm2Jae^*/J) and C57BL/6 mice heterozygous for the *Taf4b*-deficiency mutation (in exon 12 of the 15 total exons of the *Taf4b* gene that disrupts the endogenous *Taf4b* gene) were mated for collections used for mRNA isolation, meiotic spread preparation, and ovarian immunofluorescence. Timed matings were estimated to begin at day 0.5 by evidence of a copulatory plug. The sex of the embryos was identified by confirming the presence or absence of testicular cords. Genomic DNA from tails was isolated using Qiagen DNeasy Blood & Tissue Kits (Cat #: 69506) for PCR genotyping assays. All animal protocols were reviewed and approved by Brown University Institutional Animal Care and Use Committee and were performed in accordance with the National Institutes of Health Guide for the Care and Use of Laboratory Animals.

### Immunofluorescence of fetal ovaries

Prenatal ovaries were harvested at E14.5, cleaned of excess fat, and fixed in 4% formaldehyde solution for 2 hours before embedding in Optimal Cutting Temperature (OCT) compound. Ovaries were serially sectioned at 8 μm on a Thermo-Shandon Cryotome E Cryostat onto positively charged glass slides and washed twice for 5 min in PBST [1X PBS containing 0.1% Tween-20 (Fisher Scientific)] at room temperature. Tissue sections were then incubated in blocking buffer [5% goat serum (Sigma-Aldrich) in PBST] for 1 hour at room temperature. Slides were stained with rat anti-Tra98 (Abcam, ab82527) and rabbit anti-Stra8 (Abcam, ab17722) primary antibodies at 1:100 in blocking buffer and incubated overnight at 4°C on a shaker at low speed. Slides were then washed three times for 10 minutes each in PBST at room temperature. Slides were stained with goat anti-rat Alexa 488 (Abcam, ab150165) and goat-anti rabbit Alexa 555 (Abcam, ab150086) secondary antibodies at 1:500 in blocking buffer and counterstained with 4′,6-diamidino-2-phenylindole (DAPI; Vector Laboratories) and incubated for 1 h at room temperature. Slides were washed three times for 10 minutes each in PBST at room temperature before mounting coverslips with Vectashield Antifade Mounting Medium (Vector Laboratories). A secondary antibody only control was included to compare background staining. Images were taken at 20X magnification on a Zeiss Axio Imager M1 microscope (Carl Zeiss).

All images were analyzed using FIJI, and germ cell densities were determined by counting the number of Tra98 or Stra8-positive cells per section and dividing the number by the DAPI area of the ovary to obtain cells/mm^2^ (Schindelin et al., 2012). The cell counter plugin (Plugins>Analyze>Cell Counter) was used to annotate Tra98 or Stra8-positive cells. The freehand tool was used to outline the tissue section and analyze measurements (Analyze>Measure) was used to determine the area within the outline. For each genotype, three spatially dispersed sections from each ovary obtained from three mice were quantified (3 mice per genotype x 2 ovaries per mouse x 3 sections per ovary = 18 sections per genotype). Results were averaged and significance determined by an unpaired two-tailed T-test. For all graphs, dots represent individual sections counted, bar height represents the sample mean, and error bars represent standard error of the mean. All statistical analyses and graphs were generated using Graphpad/Prism version 10.1.1.

### Preparation of meiotic prophase I spreads

Ovaries were harvested at E16.5, E18.5, or PND0 and put in PBS prewarmed at 37°C until their use for spread preparation. Ovaries were then incubated in hypotonic extraction buffer [30 mM Tris, 50 mM sucrose, 17 mM trisodium citrate dihydrate, 5 mM EDTA, 0.5 mM DTT, and 0.5 mM phenylmethylsulphonyl fluoride (PMSF), pH 8.2] for 30 minutes at 37°C. After HEB incubation, ovaries were gently teased apart with 30-gauge hypodermic needles in 100 mM sucrose at room temperature. The single cell suspension was then pipetted onto slides wetted in 1% PFA with 0.2% Triton X-100 and allowed to settle overnight in a humid chamber at 37°C. The next day, slides were air-dried, at room temperature for approximately 2-3 hours. Slides were washed 2X for 5 min each in 0.4% Photo-Flo (Kodak) in PBS and then 1X for 5 min in 0.4% Photo-Flo in sterile water. After washes, slides were air-dried again for approximately one hour and then stored at -80°C.

### Immunofluorescence of meiotic prophase I spreads

After storage at -80°C, spreads were brought to room temperature by washing in PBST [1X PBS containing 0.1% Tween-20 (Fisher Scientific)] for 15 mins. Slides were then incubated in blocking buffer [5% goat serum (Sigma-Aldrich) in PBST] for 45 mins at room temperature. Slides were stained with primary antibodies diluted in blocking buffer and incubated overnight at room temperature. Primary antibodies used were mouse SYCP3 (Santa Cruz, sc74659, 1:100 dilution), rabbit SYCP3 (Abcam, ab15093, 1:100 dilution), mouse γH2AX (Millipore, 05-636. 1:100 dilution), rabbit RAD51 (Abcam, ab176458, 1:250 dilution), and rabbit TAF4b (as previously described, Grive et al. 2016, 1:100 dilution). Slides were then washed three times for 5 mins each in PBST at room temperature. Slides were stained with secondary antibodies at a 1:500 dilution in blocking buffer for 1 hr at room temperature and counterstained with 4′,6-diamidino-2-phenylindole (DAPI; Vector Laboratories). Secondary antibodies used were either goat anti-rabbit Alexa 488 (Abcam, ab150081), goat anti-mouse Alexa 488 (Abcam, ab150077), goat anti-rabbit 555 (Abcam, ab150086), or goat anti-mouse 555 (Abcam, ab150118). Slides were washed three times for 5 mins each in PBST at room temperature before mounting coverslips with Vectashield Antifade Mounting Medium (Vector Laboratories). A secondary antibody only control was included to compare background staining.

Images were taken at 100X magnification on a Zeiss Axio Imager M1 microscope (Carl Zeiss), and all image analyses were performed using FIJI. Images of 50-100 spreads were isolated from each slide, and each image was cropped to a 30–50 μm square to ensure the area only contained an individual chromatin spread. All analyses were performed on images in .czi file format. Representative images were saved in .tiff format to improve accessibility while maintaining raw data integrity. All image montages shown were generated by first converting the .czi image to an RGB (Image>Type>RGB Color) and then converting the RGB to a montage using the RGB to Montage plugin with a greyscale colorization and 10 μm scalebar.

For data shown in **Figure 4**, SYCP3 configuration was used to determine prophase I substage. For each timepoint, the total number of spreads analyzed from three animals per genotype were pooled to compare the percentage of spreads in each stage between *Taf4b*+/+ and *Taf4b*-/-genotypes. A Chi-square test was performed in Graphpad/Prism (v 10.1.1) to determine statistical significance of the difference in prophase I substage distribution between genotypes.

A previously published FIJI macro script was used for quantification of fluorescence intensity shown in **Figure 2** and **Figure 5** (Alexander et al., 2023). Images used for input into the macro were in .czi format with DAPI in blue, SYCP3 in green, and either TAF4b or γH2AX in red. Intensity of DAPI and either TAF4b or γH2AX was measured using Otsu’s thresholding. Intensity of TAF4b or γH2AX was normalized to DAPI to account for variations in DNA content amongst spreads. A copy of the FIJI macro used can be found in Alexander et al. (2023). Fluorescence intensity quantification was performed on all images, and then spreads were grouped based on prophase I substage. An ordinary ANOVA with multiple comparisons was used to compare TAF4b intensity amongst substage groups (**Figure 2A**). An unpaired two tailed t-test was used to compare intensity between substage groups (**Figure 2B**) and genotypes at each substage (**Figure 5**). For all graphs, dots represent individual spreads measured, bar height represents the sample mean, and error bars represent standard error of the mean. All statistical analyses and graphs were generated using Graphpad/Prism version 10.1.1. For data shown in **Figure S3**, RAD51 foci were counted manually for each spread. Spreads were then grouped based on prophase I substage, and RAD51 foci counts were compared between *Taf4b+/+* and *Taf4b-/-*. An unpaired two tailed t-test was used to compare foci numbers between genotypes at each substage. For all graphs, dots represent individual spreads measured, bar height represents the sample mean, and error bars represent standard error of the mean. All statistical analyses and graphs were generated using Graphpad/Prism version 10.1.1.

### Embryonic ovary dissociation and fluorescence-activated cell sorting

To dissociate ovarian tissue into a single-cell suspension, embryonic ovaries were harvested and placed in 0.25% Trypsin/EDTA and incubated at 37°C for approximately 45 minutes for E18.5 as previously described (Gura et al., 2020; Gura et al., 2022). Eppendorf tubes were subject to vigorous pipetting at 15 minute intervals to dissociate tissue throughout incubation. Trypsin was neutralized with FBS. Cells were pelleted at 1,500 RPM for 5 minutes, the supernatant was removed, and cells were resuspended in 100 μL PBS. The cell suspension was strained through a 35 μm mesh cap into a FACS tube (Gibco REF # 352235). Propidium iodide (ThermoFisher #P3566) was added at a 1:100 dilution to the cell suspension as a live/dead distinguishing stain. Fluorescence-activated cell sorting (FACS) was performed using a Becton Dickinson FACSAria III in the Flow Cytometry and Cell Sorting Core Facility at Brown University. A negative control non-GFP^+^ mouse tissue was used for each experiment to establish an appropriate GFP signal baseline. Dead cells were discarded and the remaining cells were sorted into GFP^+^ and GFP^-^ samples in PBS at 4°C for each embryo.

For RNA-seq analysis, GFP^+^ cells from each individual embryo were kept in separate tubes and were then spun down at 1,500 RPM for 5 minutes and were then resuspended in 150 µL Trizol (ThermoFisher # 1556026). The number of cells for each sample sequenced can be found in **Figure S4**. Samples were stored at -80°C.

### RNA-sequencing

Embryonic germ cells resuspended in Trizol were shipped to GENEWIZ/Azenta Life Sciences LLC. (South Plainfield, NJ, USA) on dry ice. RNA Extraction, sample QC, library preparation, sequencing reactions, and initial bioinformatic analysis were conducted at GENEWIZ/Azenta Life Sciences LLC. Total RNA was extracted from cells following the Trizol Reagent User Guide (Thermo Fisher Scientific). Extracted RNA samples were quantified using Qubit 2.0 Fluorometer (Life Technologies, Carlsbad, CA, USA) and RNA integrity was checked using Agilent TapeStation 4200 (Agilent Technologies, Palo Alto, CA, USA).

SMARTSeq HT Ultra Low Input Kit was used for full-length cDNA synthesis and amplification (Clontech, Mountain View, CA), and Illumina Nextera XT library was used for sequencing library preparation. Briefly, cDNA was fragmented, and adaptor was added using Transposase, followed by limited-cycle PCR to enrich and add index to the cDNA fragments. Sequencing libraries were validated using the Agilent TapeStation, and quantified by using Qubit Fluorometer as well as by quantitative PCR (KAPA Biosystems, Wilmington, MA, USA). The sequencing libraries were multiplexed and clustered onto a flowcell on the Illumina NovaSeq instrument according to manufacturer’s instructions. The samples were sequenced using a 2x150bp Paired End (PE) configuration. Image analysis and base calling were conducted by the NovaSeq Control Software (NCS). Raw sequence data (.bcl files) generated from Illumina NovaSeq was converted into fastq files and de-multiplexed using Illumina bcl2fastq 2.20 software. One mis-match was allowed for index sequence identification.

### Bulk RNA-sequencing data analysis

All computational scripts regarding bulk RNA-seq analysis used in this publication will be available to the public. E18.5 *Taf4b+/+*, *Taf4b+/-*, and *Taf4b-/-* RNA sequencing data were generated in this study and will be available to the public in NCBI GEO. E16.5 *Taf4b+/-* and *Taf4b-/-* RNA sequencing data are available in NCBI Gene Expression Omnibus (GEO) under accession number GSE174366.

All raw fastq files were processed on Brown University’s high-performance computing cluster. Reads were quality-trimmed and had adapters removed using Trim Galore! (v 0.6.6) with the parameters –nextera -q 10. Samples before and after trimming were analyzed using FastQC (v 0.11.9) for quality and then aligned to the Ensembl GRCm38 using HiSat2 (v 2.2.1) (Pertea et al., 2016). Resulting sam files were converted to bam files using Samtools (v 1.16.1) (Li et al., 2009).

To obtain TPMs for each sample, StringTie (v 2.2.1) was used with the optional parameters -A and -e. A gtf file for each sample was downloaded and, using RStudio (R v 4.3.2), TPMs of all samples were aggregated into one comma separated (csv) file using a custom R script. To create interactive Microsoft Excel files for exploring the TPMs of each dataset the csv of aggregated TPMs was saved as an Excel spreadsheet, colored tabs were added to set up different comparisons, and a flexible Excel function was created to adjust to gene name inputs. To explore the Excel files, click on the ‘TPM_Quick_Calc’ tab highlighted in yellow, and type the gene name of interest into the highlighted yellow boxes (**Tables S1 and S2**).

To obtain count tables, featurecounts (Subread v 2.0.2) was used (Liao et al., 2014). Metadata files for dataset were created manually in Excel and saved as a csv file. These count tables were used to create PCA plots by variance-stabilizing transformation (vst) of the data in DESeq2 (v 1.42.0) and plotting by ggplot2 (v 3.5.0) (Love et al., 2014). DESeq2 was also used for differential gene expression analysis, with count tables and metadata files used as input. We accounted for potential litter effects in our mouse oocytes by setting it as a batch parameter in DESeq2. For the volcano plot, the output of DESeq2 was used and plotted using ggplot2. DEG lists were used for ClusterProfiler (v 4.10.0) input to create dotplots of significantly enriched GO categories for all DEGs, downregulated DEGs and upregulated DEGs. Heatmaps of curated gene lists were generated by using Complex Heatmap (v 2.18.0) using a z-score matrix of normalized counts generated by DESeq2 as input.

### Single cell RNA-sequencing data analysis

Samples from GSE136441, GSE128553, GSE130212, and GSE134339 were downloaded from the NCBI GEO onto Brown University’s high-performance computing cluster at the Center for Computation and Visualization. The fastq files from each dataset were aligned using Cell Ranger (v 6.0.0) count and then aggregated using Cell Ranger aggr. The resulting output from aggr was used as input for Seurat (v 3.9.9) in RStudio (R v 4.0.2) (Stuart et al., 2019). Seurat was used to select for high-quality high quality cells based on the following parameters: nFeature_RNA > 1000, nCount_RNA < 40000, and percent.mt < 20. These data were then passed to Monocle3 (v 0.2.3) for pseudotime analysis and generating uniform manifold approximation and projection (UMAP) and gene expression data (Cao et al., 2019; Qiu et al., 2017; Trapnell et al., 2014).

## Acknowledgements

We thank Dr. Alison Delong, Dr. Ashley Webb, Dr. Karen Schindler, Dr. Kathryn Grive, and Dr. Robbert Creton for their helpful input on these studies. We thank the Center for Computation and Visualization (CCV) at Brown University for providing computational resources needed to complete scRNA-seq and RNA-seq data analysis. We are especially grateful to CCV team members Ashok Ragavendran, Joselynn Wallace, Eric Salomaki, August Guang, Paul Cao, Jordan Lawson, Prasad Bandarkar, and Prithvi Thakur for their advice and support. We thank Dr. Adriana Alexander for their expertise and advice regarding quantitative immunofluorescence of prophase I spreads. We thank Kevin Carlson and the Brown University Flow Cytometry and Sorting Facility for expertise completing the flow sorting of Oct4-EGFP gonads. The Brown University Flow Cytometry and Sorting Facility has received generous support in part by the National Institutes of Health (NCRR Grant No. 1S10RR021051) and the Division of Biology and Medicine, Brown University. As much of our insights were gained by reprocessing publicly available datasets, we greatly appreciate both the researchers that generated and shared the data initially and the respective repositories for making them available. We are grateful to the NICHD/NIH for their generous support through awards 1F31HD097933, 1F31HD105340, MAG and KMA, respectively and 1R01HD091848 and 1R01HD113567 to RNF. We also thank the US-Israel Binational Science Foundation (BSF) for their support to RNF (2019285).

---

**Figure S1.**
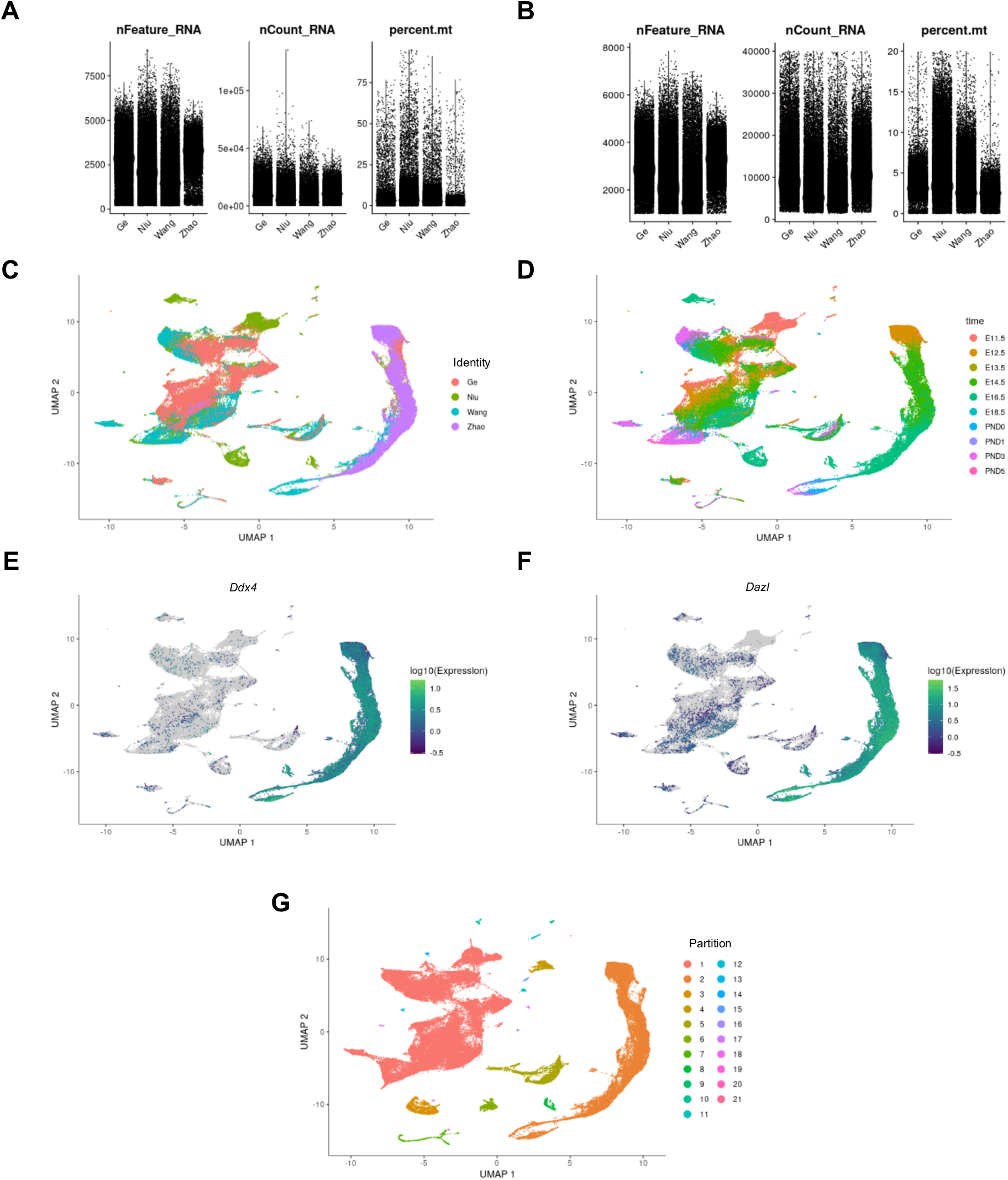
Overview scRNA-seq dataset integration. Scattered boxplots showing the QC metric distributions of the four individual datasets before (A) and after (B) setting thresholds for number of features (nFeature), number of counts (nCount), and percent mitochondrial content (percent.mt). UMAP of all cells that passed QC colored by dataset origin (C) timepoint (D) log10 expression of germ cell markers *Ddx4* (E) or *Dazl* (F) and unbiased partition assignment (G).

**Figure S2.**
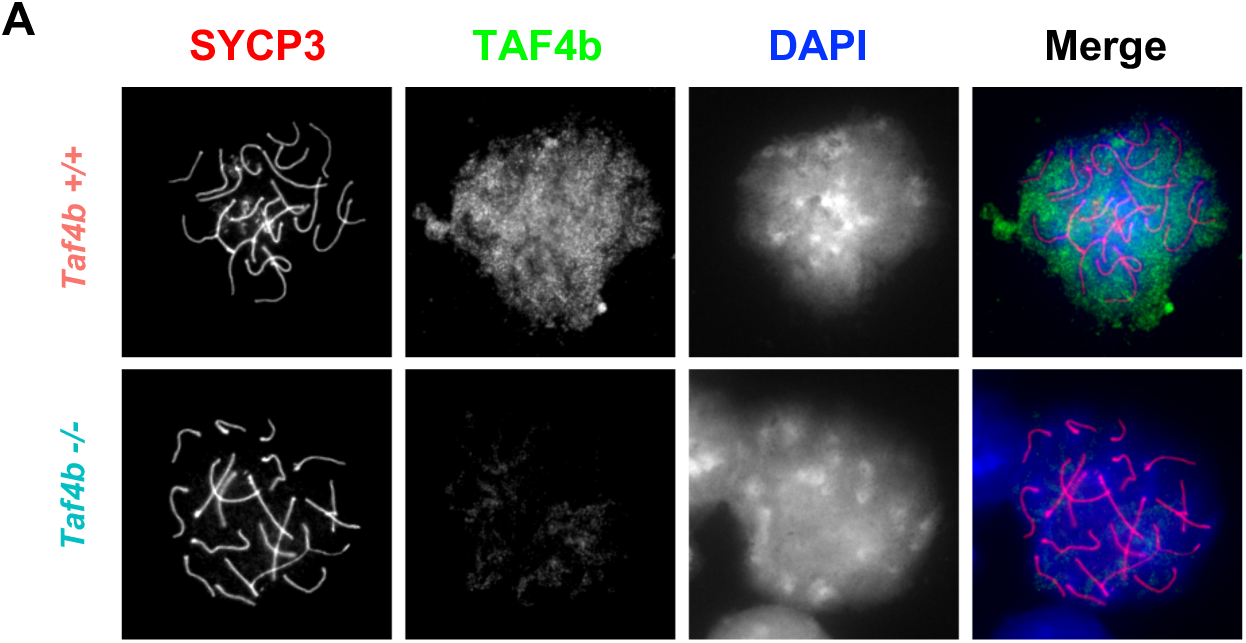
Validation of TAF4b antibody use in immunofluorescence. (A) Images of pachytene chromatin spreads from *Taf4b*+/+ (top) and *Taf4b*-/- (bottom) E18.5 ovaries stained for TAF4b (green), SYCP3 (red) and DAPI (blue).

**Figure S3.**
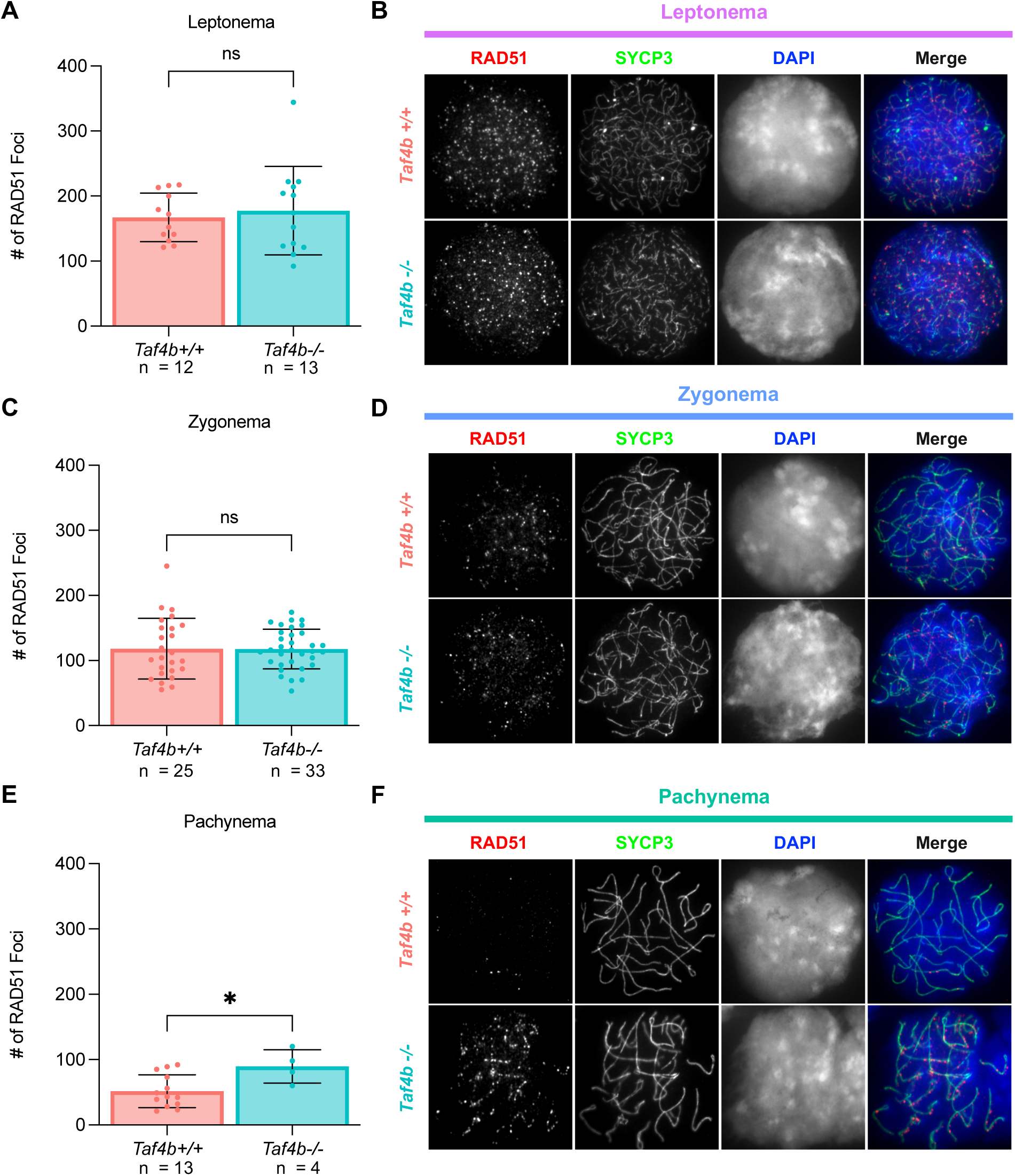
*Taf4b-/-* oocytes have elevated levels of RAD51 foci during pachynema. Quantification of RAD51 foci in chromatin spreads during leptonema (A) zygonema (C) and pachynema (E). Spreads were pooled from two E16.5 mice per genotype, n = the number of spreads analyzed. Images of spreads stained with RAD51 (red) SYCP3 (green) and DAPI (blue) from each genotype during leptonema (B) zygonema (D) and pachynema (F). Dots represent individual spreads, bar height represents the sample mean, and error bars represent standard error of the mean. Statistical significance for stage specific comparisons was determined using a two-tailed T-test, ns=not significant, and * p <0.05.

**Figure S4.**
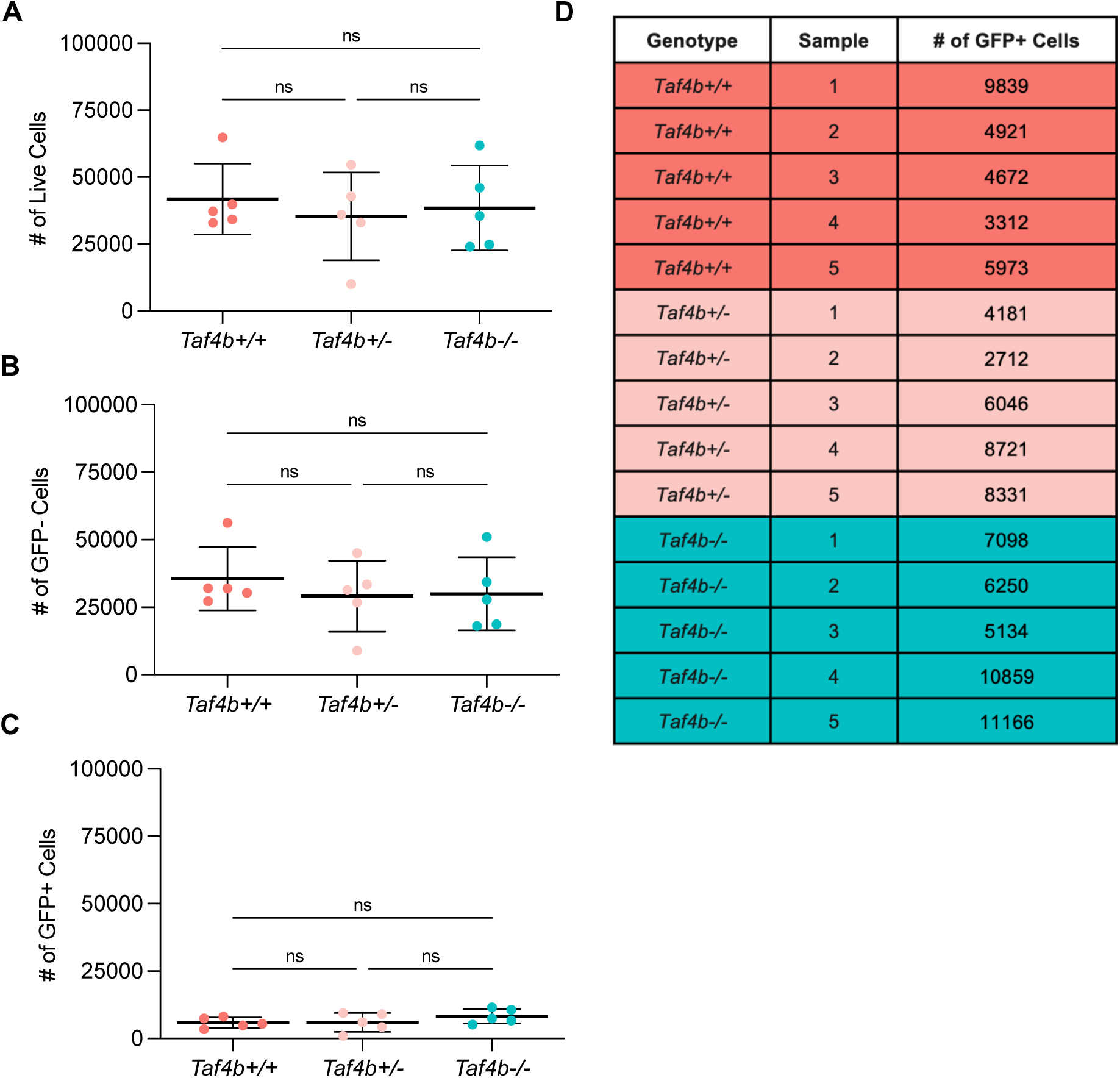
FACS summary of E18.5 samples sent for bulk RNA-seq. Charts depicting the number of live (A) GFP^-^ (B) and GFP^+^ (C) cells collected per sample during FACS of E18.5 *Taf4b+/+, +/-*, or *-/-*;Oct4-EGFP ovaries. Dots represent an individual sample, bars represent sample mean, and error bars represent standard deviation. Statistical significance was determined using an ordinary one-way ANOVA with multiple comparisons, ns = not significant. (D) Table of GFP^+^ cell numbers per sample submitted for bulk RNA-sequencing with Azenta/Genewiz.

**Figure S5.**
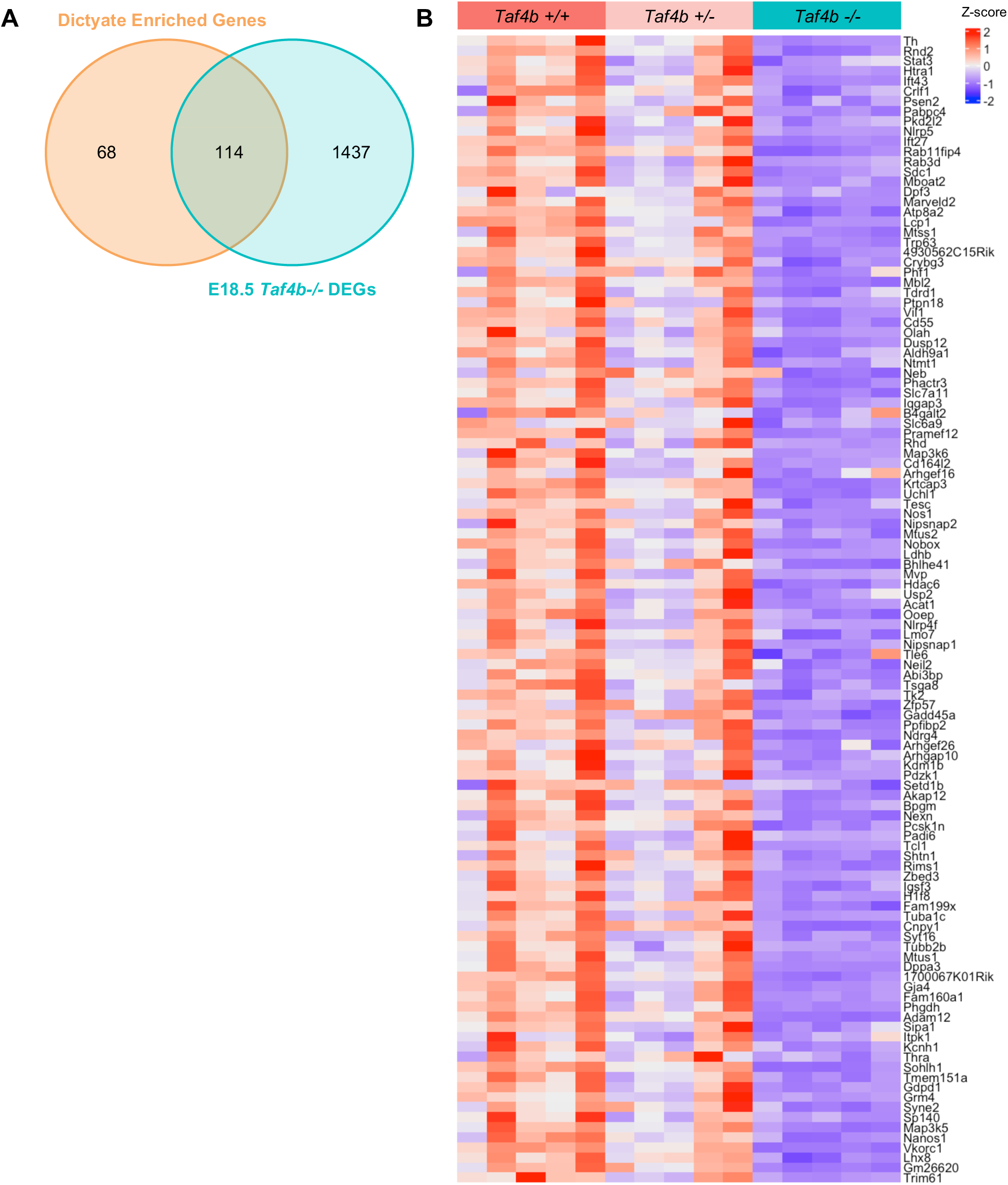
Reduced expression of dictyate enriched genes in E18.5 *Taf4b-/-* oocytes. (A) Venn diagram of E18.5 *Taf4b-/-* DEG list compared with dictyate enriched gene list (Niu & Spradling, 2022). (B) Heatmap of 114 dictyate enriched genes that are also differentially expressed in E18.5 *Taf4b-/-* oocytes.

**Figure S6.**
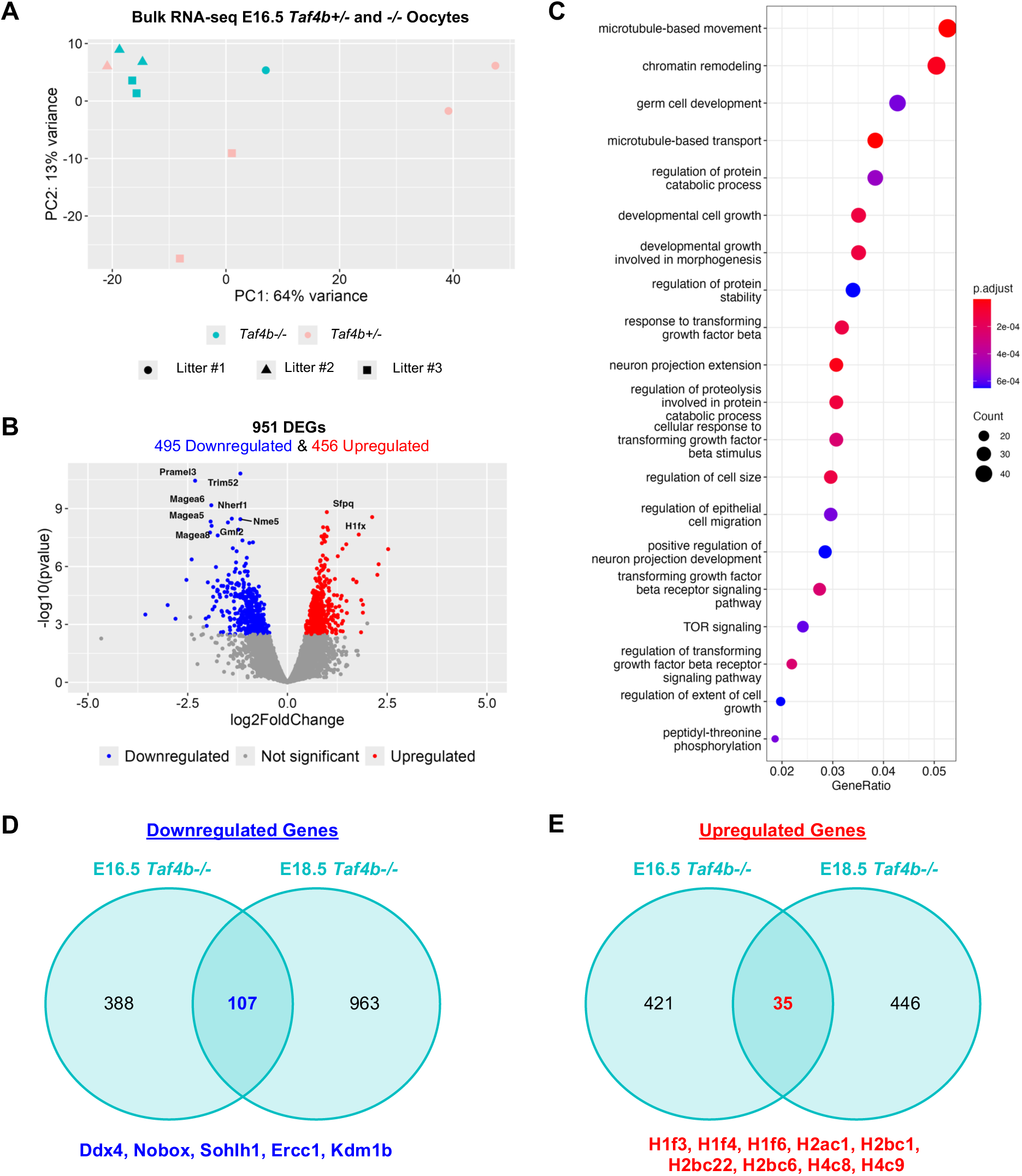
Comparison of *Taf4b-/-* DEGs at E16.5 and E18.5. (A) PCA plot of E16.5 samples labelled based on genotype and collection date. (B) Volcano plot of DEGs (protein-coding, padj <0.05, avg TPM > 1) with top 10 most significant labelled. (C) Dotplot of GO biological process analysis of 951 DEGs. Venn diagram of genes that were downregulated (D) or upregulated (E) *Taf4b-/-* oocytes at E16.5 and E18.5.

**Figure S7.**
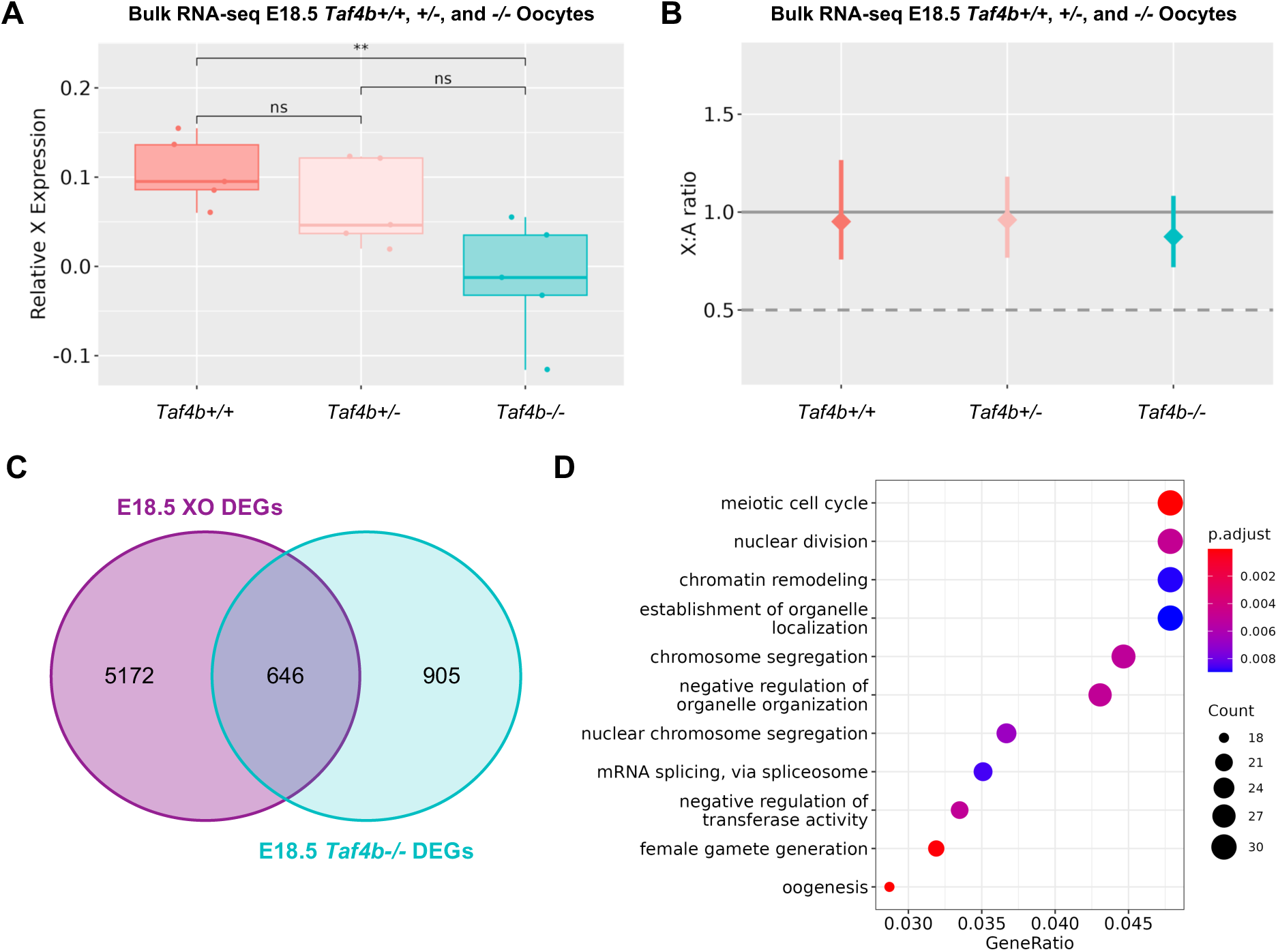
Reduced X chromosome expression in E18.5 *Taf4b-/-* oocytes. (A) Box plots of relative X expression (RXE) calculations after filtering for avg TPM > 1 and adding pseudocounts for log transformation of *Taf4b*+/+, *+/-*, and *-/-* samples. Statistical significance was determined using Welch’s T-test, ** p <0.01. Boxplot shows interquartile range from 25^th^ to 75^th^ percentile and median (solid line), whiskers represent the minimum and maximum values, and dots represent individual samples. (B) X:A ratio plot comparing *Taf4b*+/+, *+/-*, and *-/-* samples based on pairwise CI calculations performed after filtering for avg TPM > 1. Solid gray line represents full dosage compensation between X chromosomes and autosomes. Dashed gray line represents half dosage compensation of the X chromosome with the autosomes. Median as well as upper and lowers confidence intervals are plotted. (C) Venn diagram of E18.5 *Taf4b-/-* DEG list compared with E18.5 XO DEGs (protein-coding, padj <0.05, avg TPM > 1) published previously (Gura et al., 2022) (D) Dotplot of GO biological process analysis of the 642 DEGs shared between E18.5 *Taf4b-/-* and E18.5 XO oocytes.

